# Competition delays multi-drug resistance evolution during combination therapy

**DOI:** 10.1101/2020.05.27.119537

**Authors:** Ernesto Berríos-Caro, Danna R. Gifford, Tobias Galla

## Abstract

Combination therapies have shown remarkable success in preventing the evolution of resistance to multiple drugs, including HIV, tuberculosis, and cancer. Nevertheless, the rise in drug resistance still remains an important challenge. The capability to accurately predict the emergence of resistance, either to one or multiple drugs, may help to improve treatment options. Existing theoretical approaches often focus on exponential growth laws, which may not be realistic when scarce resources and competition limit growth. In this work, we study the emergence of single and double drug resistance in a model of combination therapy of two drugs. The model describes a sensitive strain, two types of single-resistant strains, and a double-resistant strain. We compare the probability that resistance emerges for three growth laws: exponential growth, logistic growth without competition between strains, and logistic growth with competition between strains. Using mathematical estimates and numerical simulations, we show that between-strain competition only affects the emergence of single resistance when resources are scarce. In contrast, the probability of double resistance is affected by between-strain competition over a wider space of resource availability. This indicates that competition between different resistant strains may be pertinent to identifying strategies for suppressing drug resistance, and that exponential models may overestimate the emergence of resistance to multiple drugs. A by-product of our work is an efficient strategy to evaluate probabilities of single and double resistance in models with multiple sequential mutations. This may be useful for a range of other problems in which the probability of resistance is of interest.

## 1. Introduction

The rise of drug resistance has triggered studies into different treatment regimes, aimed to prevent or delay the emergence of resistance. Combination therapies have attracted special attention, due to their effectiveness against viral, bacterial, and fungal infections, as well in the context of cancer (see e.g. Devita Jr et al. (1975); Bonhoeffer et al. (1997); Livermore (2005); Baym et al. (2016)). Combination therapies have also shown success in managing HIV (Richman (2001)), malaria (Nosten and White (2007)), and tuberculosis (Mitchison and Davies (2012)).

The ability of combination therapies to counteract antibiotic resistance relies on the idea that simultaneous acquisition of resistance to multiple antibiotics is extremely rare. For independent resistance mutations, this occurs with a rate equal to the product of mutation rates for resistance to each drug. For bacteria these are in the range of approximately 10^−6^ to 10^−10^ per genome replication (see Krašovec et al. (2017)). Multi-resistance is therefore more likely to emerge via sequentially acquisition of resistance to each drug Bonhoeffer et al. (1997). This is what combination therapies aim to prevent. These therapies can have limitations, however, for example due to differences between drugs in how quickly they are absorbed (i.e. pharmacokinetics, see Yeh et al. (2006); Peña-Miller et al. (2013)).

Having the ability to predict the emergence of single or double resistance from mathematical or computational models may help to develop strategies to reduce resistance. Since mutations occur as random events, one of the main aims is to compute or estimate the probability that resistant cells are present in the population at a certain time after the drug treatment has started. Several theoretical approaches have been proposed, with a particular focus on drug treatments in cancer cells (see Altrock et al. (2015) for an overview). Other modelling work has focused on resistance in viral dynamics (see e.g. Alexander and Bonhoeffer (2012)). A systematic review of models of antimicrobial resistance can be found in Niewiadomska et al. (2019).

Michor *et al.* (Michor et al. (2006)) estimated the probability of extinction of a branching process with multiple types of mutations, for populations consisting entirely of sensitive cells at the start of the treatment. Their approach allows one to obtain the probability of successful therapy, that is, the therapy that kills both sensitive and mutant cancer strains at very long times after the treatment started. The calculations are based on the work of Iwasa and co-authors (see Iwasa et al. (2003, 2004)), where extinction probabilities were derived for an exponential growth model from a generating-function approach. Similarly, Iwasa et al. (2006), estimated the probability of single resistance at the point in time at which the total cell population reaches a certain size. This was extended in Haeno et al. (2007) to a case with single and double resistants. This latter approach succeeded in obtaining the probability of having at least one double mutant before the total population has reached a particular size.

Later, Foo and Michor (2010) proposed a simpler approach, based on using the extinction probability of a single-type birth-death process, to estimate the emergence of single resistance in dosing schedules that affect both birth and death rates. Recently, the effects of pharmacokinetics were studied by Chakrabarti and Michor (2017). Related work can also be found in Bozic et al. (2013) for a process with multiple types of resistance, see also Alexander and MacLean (2019) for an experimental approach.

The work mentioned so far focuses largely on models with exponential growth, effectively ignoring interactions between cells. This is an approximation, but it works well for the early stages growth when non-linear interaction effects are not relevant (see e.g. Monod (1949)). In particular, this assumption is valid when the population size has not yet saturated at carrying capacity.

Although exponential growth can serve as a good approximation to study the emergence of single mutations, it may not be appropriate when multiple mutations take place in sequence. This is because the later mutations can occur during advanced phases of growth. At that stage interactions between cells may have become relevant, in particular when the population approaches its carrying capacity. In order to model such instances, one needs to go beyond simple unconstrained reproduction. The purpose of this work is to show how the choice of growth model affects predictions for the emergence of single and double resistance under combination therapy. We describe the evolution of resistance by means of three stochastic models: (i) A model with no interactions between cells, leading to exponential growth if growth rates are constant; (ii) A model in which each strain follows a logistic growth law, but where there are no interactions between the different strains; (iii) Logistic growth with competition between strains for a common resource. For each of the model we consider constant and time-dependent per capita birth and death rates. We derive analytical estimates for the probability of single and double resistance, and compare these with numerical simulations. The formalism to predict the emergence of resistance builds upon the work in Foo and Michor (2010).

Our results show that the prediction of single resistance in the logistic growth models is different from that for the model with exponential growth when the availability of resources is low. The probability of double resistance varies across the different growth models for a larger range of parameters (such as birth rates of sensitive or resistant strains, or the initial cell number). This difference to the exponential model is more pronounced in the model with competition between different strains. The growth of source strains for double mutants may then saturate before double mutants appear. This earlier saturation can alter the optimum treatment strategy, i.e., the therapy that maximises the time at which first double mutants emerge.

The remainder of this paper is set out as follows. In Section 2 we provide the mathematical background definitions for our study. In particular, we describe the different growth models and we define the model we use for the effects of treatments on these growth laws. Section 3 contains our main analytical results. We derive approximations for the probability of single and double resistance, and describe how we evaluate the resulting numerical expressions efficiently. In Section 4 we then discuss the these predictions for models with constant parameters. Time-dependent dosing protocols are studied in Section 5. In Section 6 we discuss limitations of our approach, before we conclude in Section 7. Further details of our calculations, and additional results can be found in the Supplementary Material.

## 2. Stochastic Model

### 2.1. General definitions

We focus on a cell population subjected to combination therapy of two different types of drugs labeled 𝒜 and ℬ respectively. The drugs may act concurrently, depending on the dosing schedule. Cells can develop resistance to one drug (single resistance) or to both drugs (double resistance) via independent mutations. Double resistance is acquired sequentially, that is, after acquiring first single resistance to either of the two drugs. We will not consider the possibility of acquiring resistance to both drugs through a single mutation, as this is sufficiently rare (see e.g. Bonhoeffer et al. (1997)). Our model excludes back mutations as these are also very unlikely (see e.g. Allen et al. (2017)).

To model how the strains acquire resistance, we consider a multi-strain continuous-time birth-death process with mutations. The birth and death rates can vary over time, making the dynamics a so-called ‘non-homogeneous’ process (Bailey (1990)). There are four different strains in the population, labeled *S, A, B*, and *D*. Strain *S* denotes sensitive cells, i.e., this strain does not exhibit resistance to either drug. Cells of strain *A* are resistant to drug 𝒜, but not to ℬ, and vice versa for *B*. Strain *D* consists of double-resistant cells.

The process is described by random variables *n*_*S*_(*t*), *n*_*A*_(*t*), *n*_*B*_(*t*), and *n*_*D*_(*t*), which represent the cell numbers (or number of individuals) for each strain at time *t*. We write **n** = (*n*_*S*_, *n*_*A*_, *n*_*B*_, *n*_*D*_). Members of different strains proliferate and die with rates *b*_*i*_(*t*) and *d*_*i*_(*t*) respectively, where *i* ∈ {*S, A, B, D*}. These rates can be explicit functions of time, reflecting time-dependent treatment strategies. In birth events a single individual of the population reproduces, and in death events a single individual is removed from the population.

In birth events, one of the two types of mutations can occur. For example, an offspring of strain *S* will be of type *A* with probability *µ*_*A*_, and of type *B* with probability *µ*_*B*_. This is described by the events *S* → *S* + *A* or *S* → *S* + *B*, respectively. With the remaining probability, 1 − *µ*_*A*_ − *µ*_*B*_, the offspring is of type *S* (*S* → *S* + *S*). Similarly, an offspring of a parent type *A* is of type *D* with probability *µ*_*B*_(*A* → *A* + *D*), and an offspring of an individual of type *B* is of type *D* with probability *µ*_*A*_(*B* → *B* + *D*). The per capita birth rate *b*_*i*_ has an explicit dependence on *n*_*i*_ in the model with logistic growth without competition between strains. Each *b*_*i*_ depends on all components of **n** in the model with competition between strains. This will be detailed further below. The rates associated with the possible events are summarised in Table 1. The last column of the table indicates how entries of the state vector (*n*_*S*_, *n*_*A*_, *n*_*B*_, *n*_*D*_) change in the different types of events.

**Table 1.** Summary of the different birth and death events in the population, along with the associated rates per individual, and the total rate for any type of event in the population. The symbol 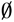 represents removal of an individual from the population (i.e., death). The change of the state vector **n**(*t*) = (*n*_*S*_(*t*), *n*_*A*_(*t*), *n*_*B*_(*t*), *n*_*D*_(*t*)) in each of the different possible events is shown in the last column.

| Transition | Rate per individual | Total event rate | Change of $\mathbf{n}(t)$ |
| --- | --- | --- | --- |
| $S \rightarrow S + S$ | $b_S(t)(1 - \mu_A - \mu_B)$ | $n_S(t)b_S(t)(1 - \mu_A - \mu_B)$ | $(1, 0, 0, 0)$ |
| $S \rightarrow \emptyset$ | $d_S(t)$ | $n_S(t)d_S(t)$ | $(-1, 0, 0, 0)$ |
| $S \rightarrow S + A$ | $b_S(t)\mu_A$ | $n_S(t)b_S(t)\mu_A$ | $(0, 1, 0, 0)$ |
| $S \rightarrow S + B$ | $b_S(t)\mu_B$ | $n_S(t)b_S(t)\mu_B$ | $(0, 0, 1, 0)$ |
| $A \rightarrow A + A$ | $b_A(t)(1 - \mu_B)$ | $n_A(t)b_A(t)(1 - \mu_B)$ | $(0, 1, 0, 0)$ |
| $A \rightarrow \emptyset$ | $d_A(t)$ | $n_A(t)d_A(t)$ | $(0, -1, 0, 0)$ |
| $A \rightarrow A + D$ | $b_A(t)\mu_B$ | $n_A(t)b_A(t)\mu_B$ | $(0, 0, 0, 1)$ |
| $B \rightarrow B + B$ | $b_B(t)(1 - \mu_A)$ | $n_B(t)b_B(t)(1 - \mu_A)$ | $(0, 0, 1, 0)$ |
| $B \rightarrow \emptyset$ | $d_B(t)$ | $n_B(t)d_B(t)$ | $(0, 0, -1, 0)$ |
| $B \rightarrow B + D$ | $b_B(t)\mu_A$ | $n_B(t)b_B(t)\mu_A$ | $(0, 0, 0, 1)$ |
| $D \rightarrow D + D$ | $b_D(t)$ | $n_D(t)b_D(t)$ | $(0, 0, 0, 1)$ |
| $D \rightarrow \emptyset$ | $d_D(t)$ | $n_D(t)d_D(t)$ | $(0, 0, 0, -1)$ |

### 2.2. Mean growth laws and production rates of mutants

The quantities *n*_*S*_(*t*), *n*_*A*_(*t*), *n*_*B*_(*t*), and *n*_*D*_(*t*) above are random variables, and differ from realisation to realisation of the stochastic birth-death dynamics. We write 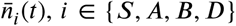 for the average cell numbers at time *t* across realisations. In the limit of infinite populations these follow the ordinary differential equations 

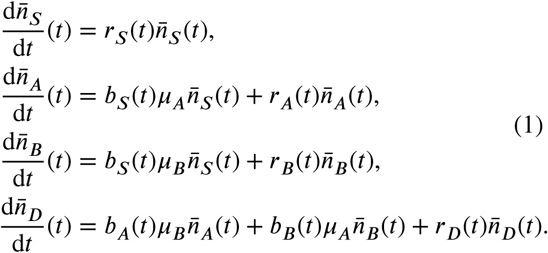

The quantities 

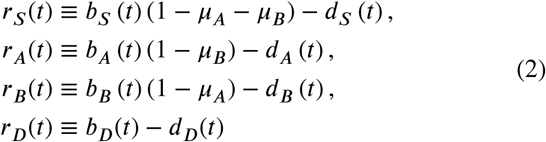

represent the net growth rates for the different strains, and result from the balance of birth and death. In the exponential growth model the *b*_*i*_(*t*) do not depend on 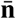. In the logistic model without competition between strains *b*_*i*_ is of the form 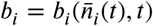 in the deterministic limit. In the model with competition between strains, finally, we have 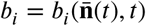. This is obtained from factorising averages in the deterministic limit, e.g.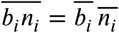, and using 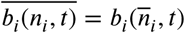. In Eq. (2) we have simply written *b*_*i*_ = *b*_*i*_(*t*) to keep the notation compact.

To proceed we will assume that the probabilities *µ*_*A*_ and *µ*_*B*_ are small compared to one, values below 10^−2^ can serve as a reference for real-world biological systems, see e.g. Krašovec et al. (2017). We use this to approximate the terms 1− *µ*_*A*_, 1− *µ*_*B*_, and 1− *µ*_*A*_ − *µ*_*B*_ in Eq. (2) by a value of unity. This means that we overestimate the number of non-mutant offspring by a small fraction. In related models it has been shown that this simplification does not significantly alter the outcome (Foo and Michor (2010)). With this simplification, the net growth rates in Eq. (2) become *r*_*i*_ = *b*_*i*_ − *d*_*i*_. The mutation rates are still present in Eqs. (1), in the terms associated with mutations from source strains (the first term in each of the growth laws for strains *A, B* and *D*).

The rates with which single-resistants of type *A* or *B* are produced by mutations are given by the total rates of the events *S* → *S* + *A* and *S* → *S* + *B*, respectively. In the fully stochastic model these are random quantities themselves, given by *W*_*A*_(*t*) = *b*_*S*_(*t*)*µ*_*A*_*n*_*S*_(*t*) and *W*_*B*_(*t*) = *b*_*S*_(*t*)*µ n*_*S*_(*t*) (see Table 1). Within the deterministic approximation we can replace this by 

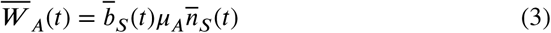

and 

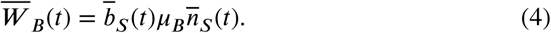

The production rate of double mutants, resulting from mutations arising in the offspring of either *A* or *B*, is the sum of the rates for the two events *A* → *A* + *D* and *B* → *B* + *D*. Focusing on mean values again, this becomes 

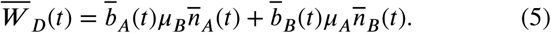

The deterministic approximation we have made effectively amounts to writing 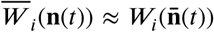. The resulting equations for the mean cell numbers are then closed in the 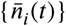, with no dependence on higher-order moments of *n*_*i*_(*t*). This approximation is only necessary in the logistic growth models defined below. For the range of parameters we explore in this paper, theoretical predictions calculated based on this simplification are typically in good agreement with numerical simulations as we will see below. We discuss limitations below in Section 6, and we show instances in which the predictions of the theory deviate from numerical simulations in Section S6 of the Supplementary Material.

### 2.3. Specific growth laws

We will explore three different scenarios. The first is exponential growth, and describes, for example, bacterial populations with unlimited resources so that growth can continue indefinitely (e.g. as it is the case in so-called ‘continuous’ culture systems with constant growth rates, see Hoskisson and Hobbs (2005)). The second scenario entails logistic growth for each strain, but with no competition between individuals of different strains. This describes situations where resources are limited, but each strain exploits a different resource (such as may occur if a resistance mutation allows a strain to occupy a new ecological or spatial niche, Moreno-Gamez et al. (2015); Feder et al. (2019)). The growth of any one strain is then limited by the number of individuals of that strain, but not by individuals of other strains. The third scenario is logistic growth with competition within and between strains. The different strains compete for the same resources, such that the growth of any one strain is limited by the presence of all strains. This leads to competitive Lotka-Volterra equations (Schoener (1976)). We provide detailed mathematical definitions for each of the three scenarios below.

Throughout this paper and for every type of growth law, the initial population at time *t* = 0 is assumed to consist of *n*_0_ sensitive cells and no resistant mutants, i.e., the initial condition is **n**(*t* = 0) = (*n*_0_, 0, 0, 0).

#### 2.3.1. Exponential growth model (EG)

In this model, the per capita birth and death rates do not depend on the cell number of any of the strains, i.e., the *b*_*i*_(*t*) and *d*_*i*_(*t*) may be time-dependent (reflecting time-varying drug concentrations), but they are not functions of **n**. The differential equations for the 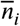 are then linear in 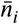. In the special case of constant per capita growth and death rates, the equations admit analytical solutions in the form of elementary exponential functions (see Section S2 of the Supplementary Material). When the rates are time-dependent, the solutions can be expressed as 

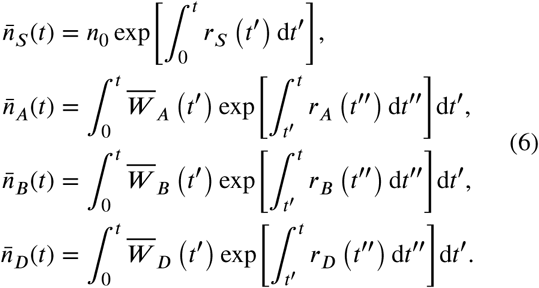

Even though these are not strictly exponential functions in the case of time-dependent rates we will nevertheless refer to this type of growth law as ‘exponential growth’, and use the shorthand ‘EG’.

It is straightforward to interpret the first relation in Eq. (6). The growth of strain *S* is described by a single-strain birth-death process with initial condition *n*_0_, and the net growth rate at time *t*′ is *r*_*S*_(*t*′) (this may be negative, in which case the number of individuals of type *S* reduces with time). For the resistant strains (*A, B* and *D*) the term 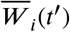 represents the (mean) rate with which individuals of this strain are produced in mutation events at time *t*′. The exponential term then accounts for their replication (or removal) between times *t*′ and *t* through a birth-death process with net reproduction rate *r*_*i*_(*t*″) at time *t*″. Thus, the integral over *t*′ counts all new mutations, and the offspring the mutant produces between the time of the mutant’s appearance, *t*′, and time *t*.

The *n*_*i*_ are not bounded in this model, implying that the cell numbers can grow to arbitrary values. This situation is not realistic as resources are limited in practice; in an infection for example, cell numbers are constrained by nutrients provided by the host (Smith and Holt, 1996; Smith, 2007). The two scenarios that we describe next hence account for limited growth.

#### 2.3.2. Logistic growth model without competition between strains (LG)

In this scenario each strain follows a logistic growth law, that is to say, the cell number grows exponentially initially, but then approaches a carrying capacity at later times. We write *k*_*i*_ for the carrying capacity of strain *i*, and assume that the net growth rate *r*_*i*_ for strain *i* depends on *n*_*i*_, but not on the cell numbers of the remaining strains. This can describe microbial communities in which the different strains consume different resources. We will use the shorthand ‘LG’ to refer to this setup (logistic growth).

There are several ways of choosing birth and death rates so that the resulting mean cell number follows a logistic growth law, see e.g. Smith and Tuckwell (1974); Parthasarathy and Krishna Kumar (1991); Goel and Richter-Dyn (2016). For example, the model could be such that the birth and death rates for strain *i* both become zero when the cell number *n*_*i*_ reaches the carrying capacity *k*_*i*_. In a stochastic model this would mean that the birth-death dynamics comes to a complete halt when the carrying capacity is reached. As a consequence *n*_*i*_ will remain fixed at *n*_*i*_ = *k*_*i*_, and no fluctuations around the carrying capacity are observed.

In our model, we make less restrictive assumptions. We only require that the cell number of a strain does not grow *on average* when it exceeds its carrying capacity (i.e. *r*_*i*_ = *b*_*i*_ − *d*_*i*_ = 0 when *n*_*i*_ ≥ *k*_*i*_ for *i* ∈ {*S, A, B, D*}). We make the choice *r*_*i*_ = 0 for *n*_*i*_ > *k*_*i*_ for reasons of consistency with the logistic growth model with competition between strains. This will be explained in more detail below. In order to allow fluctuations of the cell numbers around the carrying capacity, we choose both birth and death rates to be non-zero at and above the carrying capacity.

To specify the birth and death rates, we start from a given per capita death rate *d*_*i*_(*t*), and assume that the per capita birth rate *b*_*i*_(*t*) equals *d*_*i*_(*t*) when *n*_*i*_(*t*) *k*. This implies *r*_*i*_(*t*) = *b*_*i*_(*t*) − *d*_*i*_(*t*) = 0 for *n*_*i*_ *k*, i.e. zero net growth. The net growth for strain *i* is chosen to be positive when *n*_*i*_ < *k*_*i*_, with net growth rate *r*_*i*_(*t*) = *ρ*_*i*_(*t*)[1 − *n*_*i*_(*t*)/*k*_*i*_]. The factor in the square bracket ensures that the net growth rate reduces to zero as *n*_*i*_ approaches the carrying capacity *k*_*i*_. The quantity *ρ*_*i*_(*t*) is the intrinsic growth rate of strain *i*, describing the per capita net growth rate of the strain in the limit of small cell numbers (*n*_*i*_ ≪ *k*_*i*_). A similar approach was used in Parthasarathy and Krishna Kumar (1991) for a stochastic version of the Gompertz model. Other related works of similar models with coupled birth and death rates can be found in Matis and Kiffe (1996); Matis et al. (2003, 1998). Summarising, we use 

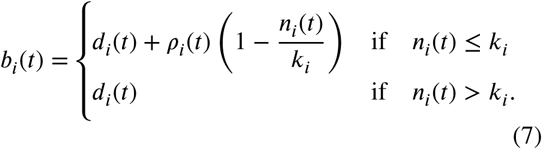

The specifics of the external influence of drug therapy are reflected in the choice of the functions *d*_*i*_(*t*) and *ρ*_*i*_(*t*). This is discussed further in Sections 5.1 and 5.2.

The dynamics discussed above only describes changes in *n*_*i*_ due to reproduction and death of strain *i*, but it does not cover production of strains *A, B* and *D* due to mutations. For example, strain *A* may have reached carrying capacity and hence there is no intrinsic net growth of this strain (*r*_*A*_ = 0), but additional individuals of type *A* are still produced when mutations occur in reproduction events of strain *S*. This is reflected by the first term on the right-hand side of the growth laws for 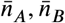 and 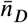 in Eq. (1). For example 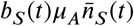 will generally be positive, even when strain *A* has reached carrying capacity. In this situation, the cell number of strain *A* fluctuates stochastically, with a net increase over time due to mutations from strain *S*. The latter occurs with a rate proportional to *µ*_*A*_, which is assumed to be much smaller than one. As a consequence mutants will be generated at a rate which is slow compared to the reproduction rate of type-*A* individuals in the growth phase before saturation.

#### 2.3.3. Logistic growth model with competition between strains (CLG)

This model is similar to the one in the previous section. Crucially though, the growth of any one strain can now be affected by the total cell number of all strains in the population. We write *n*_*T*_(*t*) = *n*_*S*_(*t*) + *n*_*A*_(*t*) + *n*_*B*_(*t*) + *n*_*D*_(*t*) for this total cell number. The model captures scenarios in which the different strains all compete for the same resource. We make the simplifying assumption that the interaction is the same between any pair of strains, such as for example pure scramble competition. There is no direct interference, cross-feeding or similar (see e.g. Jørgensen and Fath (2014)). We will use ‘CLG’ to refer to this class of growth law (‘competitive logistic growth’).

Similar to the previous section, we use the per capita birth rate 

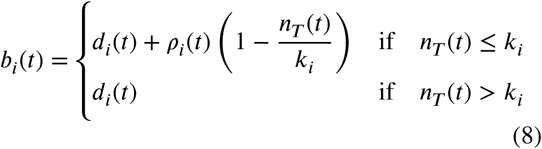

for strain *i*. The net birth-death growth rates are *r*_*i*_ = *b*_*i*_ − *d*_*i*_ as before. The term *k*_*i*_ is the carrying capacity of strain *i* in absence of its competitors.

This choice indicates that the growth rate for strain *i* will reduce to zero (*r*_*i*_ = 0) when the total cell number *n*_*T*_ exceeds *k*_*i*_. In an alternative setup with *r*_*i*_ < 0 for *n*_*T*_ > *k*_*i*_ only the strain with highest coefficient *k*_*i*_ would survive at long times. To see this, consider the case in which *n*_*T*_ > *k*_*i*_ for species *i*, but *n*_*T*_ < *k*_*j*_ for species *j*. The abundance of the first species would decline, that of the second species would continue to grow. Our approach ensures that the growth laws are such that the mean abundance 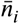 saturates when 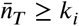. The birth and death rates for strain *i* will nevertheless remain non-zero, and hence the number of cells of type *i* will keep fluctuating in the individual-based model.

This model is based on the competitive Lotka-Volterra equations (Schoener (1976)). In their general form (see e.g. Gilpin and Ayala (1973)), these equations include an interaction matrix that accounts for the competition of any strain with any other; the interaction coefficients can – in principle – be different for different pairs of strains. We have here focused on the simple case, with all interaction coefficients set equal to unity. In previous work Gifford et al. (2019) we have used a similar setup to model growth curves obtained experimentally in populations of bacteria.

### 2.4. Effect of drug treatment on growth rates

What effects drugs have on the different strains may depend on factors such as environmental conditions, drug interactions, type of drugs used, etc. We assume synergistic effects of both drugs on the sensitive strain, i.e., the effect of the combination of both drugs on *S* is stronger than the effect of any one drug in isolation. We also assume that the growth of strain *S* will be suppressed by a larger amount in the presence of both drugs than that of any of the single-resistant strains. Growth of the double-resistant strain is not affected by any drugs in our model.

Net growth is described by the balance of birth and death rates. Depending on the type of drugs used, net growth can be affected in different ways. Bactericidal drugs (which kill bacteria) will primarily increase death rates even though birth rates may also be affected (Kohanski et al. (2010). Bacteriostatic drugs (which do not necessarily kill bacteria but slow their reproduction) will mostly affect birth rates.

In practice, however, drugs will often affect birth and death rates simultaneously. In our model drug 𝒜 reduces the birth rates of the sensitive strain *S* and of the single-mutant strain *B* (*b*_*S*_ and *b*_*B*_, respectively). Strain *A* is resistant to drug 𝒜. Similarly, drug ℬ reduces *b*_*S*_ and *b*_*A*_, but not *b*_*B*_. At the same time drug 𝒜 will also increase the death rate of strains *S* and *B* (*d*_*S*_ and *d*_*B*_, respectively), and drug ℬ increases *d*_*S*_ and *d*_*A*_. The birth and death rates for the double resistant strain (*b*_*D*_ and *d*_*D*_) are not affected by either drug.

In order to capture this scenario, we write *C*_*A*_ and *C*_*B*_ for the (dimensionless) concentrations of the two drugs, and introduce 

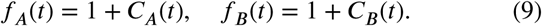

We allow drug concentrations to be a function of time to account for dosing schedules and drug pharmacokinetics. The quantities *f*_*A*_(*t*) and *f*_*A*_(*t*) describe the relative factors by which birth and death rates are reduced or enhanced, respectively. We note that *f*_*A*_ = 1 in the absence of drug 𝒜, and similarly for *f*_*B*_.

To model the effects of drugs in the exponential model we start from constant birth and death rates, labelled 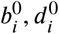. These coefficients describe the birth and death rates in the absence of any drugs (*C*_*A*_ = *C*_*B*_ = 0). The reduction of growth rates and increase of death rates is then captured as follows: 

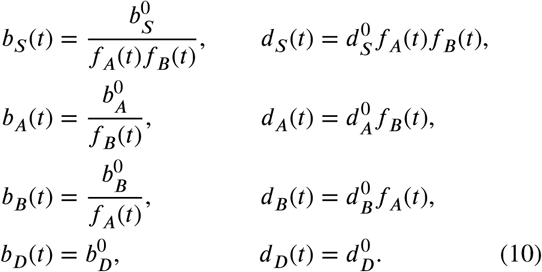

We will refer to this as an ‘exponential growth model with time-dependent drug concentrations’, even though the growth process is no longer strictly exponential when *C*_*A*_ and *C*_*B*_ are functions of time.

Equation (10) defines the *d*_*i*_(*t)* and *b*_*i*_(*t)* for the exponential model in the presence of drugs. For the logistic models with drug treatment, we use the same death rate *d*_*i*_(*t)* as in the exponential model. We then define the intrinsic growth rates *ρ*_*i*_(*t*) ≡ *b*_*i*_(*t)* − *d*_*i (*_*t*), with *b*_*i*_(*t*) and *d*_*i*_(*t*) as in Eq. (10). In this definition of *ρ*_*i*_(*t)* we use the birth rate *b*_*i*_(*t*) for the exponential model. The birth rates *b*_*i*_(*t)* for the logistic models are then constructed from *d*_*i*_(*t)* and *ρ*_*i*_(*t*) using Eqs. (7) and (8) respectively. This allows one to compare the outcome of the different growth models. When cell numbers are much smaller than the relevant carrying capacities the suppression of growth in the logistic models does not yet set in. The three models then exhibit similar behaviour.

The relations in the first line of Eq. (10) represent the reduction of the birth rate for strain *S* in the presence of any of the two drugs; similarly the death rate for strain *S* is increased. For simplicity we assume that the factors by which *b*_*S*_ is reduced are the same as the those by which *d*_*S*_ is increased. A similar approach is taken for the remaining strains. In principle, more general choices are possible, but in the spirit of constructing a stylised model capturing the essential effects, we proceed on the basis of Eq. (10).

The multiplicative setup in Eq. (10) ensures that the birth and death rates remain non-negative, even for very high drug concentrations. This is harder to ensure in an additive model, for example of the form 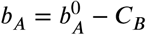.

## 3. Probability of single and double resistance

In this section, we present analytical approximations for the probability that the population has developed resistance at a given time. More specifically, we calculate the probabilities that there is at least one individual of type *A* or of type *B* in the population at time *t* (*P* [*n*_*A*_(*t*) + *n*_*B*_(*t*) > 0]), and the probability that there is at least one double-resistant *D* in the population *P* [*n*_*D*_(*t*) > 0]). The method to derive these probabilities is based on Foo and Michor (2010), where the probability of having at least one single-resistant was calculated for a linear model with exponential growth and only one type of single-resistant strain. Our contribution consists of extending this method to the non-linear growth models described in the previous section, and additionally, we also obtain the probability to find double resistance. We note that double mutants *D* can be generated from two parent sources (*A* and *B*) via mutations. We here only outline the main steps and results. Further details of the calculations can be found in the Supplementary Material (Section S1).

### 3.1. Extinction probability of single strains

The method of Foo and Michor (2010) makes use of the extinction probability in a single-species birth-death process with time-dependent birth and death rates. This is the probability that one single individual present at time *t* becomes extinct by time *T*, along with its lineage (i.e, the cell itself and all its descendants).

For strain *i*, and using the previous notation *b*_*i*_(*t*) and *d*_*i*_(*t*′) for the per capita birth and death rates at time *t*′, this probability is given by Bailey (1990); Parzen (1999) 

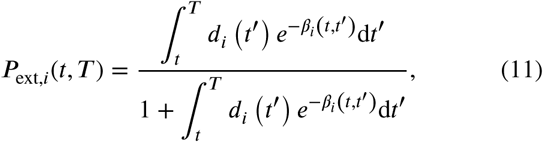

with 

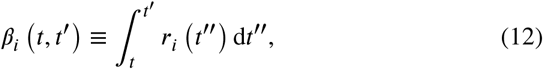

and *r*_*i*_(*t)* = *b*_*i*_(*t)* − *d*_*i*_(*t)* as before. This result is exact, provided the rates *b*_*i*_(*t)* and *d*_*i*_(*t)* are deterministic functions (for example externally determined birth and death rates). When one or both of these rates depend on the cell numbers **n** one can proceed based on an approximation in which one replaces **n** by the mean cell numbers 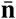. We will make use of this approximation for the logistic growth models. If *d*_*i*_ = 0, the extinction probability is zero trivially, as there is no death process. The extinction probability in Eq. (11) tends to unity for *T* → ∞ if and only if 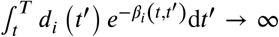. This can occur, for example, when the death rate is higher than the birth rate at all times, i.e., *r*_*i*_(*t)* is consistently negative and with it also *β*_*i*_(*t, t*′).

In order to calculate the probabilities of single or double resistance one needs to evaluate *P*_ext,*I (*_*i* ∈ {*A, B, D*}) as defined above for each growth model, for both constant and time-dependent rates. Although this can be done numerically for cases which do not admit an analytical solution, the repeated evaluation of the integrals in Eqs. (11) and (12) can become very costly. One aspect of our work is the strategy developed to carry out these integrals efficiently. This is explained below in Section 3.4, after we first describe the main steps we follow to obtain the probabilities of single and double resistance.

### 3.2. Single resistance

As described in more detail in Section S1.2.1 of the Supplementary Material, the probability of having at least one single-resistant cell of either type *A* or *B* at time *T* after the treatment has started can be expressed in the form 

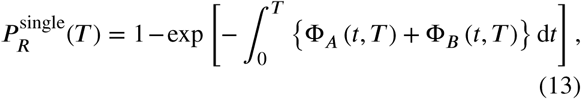

where 

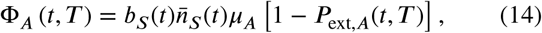

and 

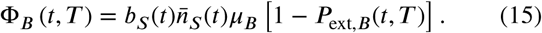

The term Φ_*i*_ *(t,T)* represents the rate with which mutants of type *I* ∈ *{A,B*} are produced at time *t* through mutations of strain *S*, requiring that the resulting mutant lineage survives until the later time *T*. We will refer to the expression in Eq. (13) as the *probability of single resistance*.

Broadly speaking the above expressions indicate that the appearance of single mutants is guaranteed (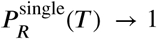 for *T* → ∞) if mutation events from the sensitive strain continue indefinitely, and if mutants produced in this way do not all die out in the long run. We would expect the latter to be the case when the growth rate *r*_*i*_(*t)* remains positive throughout for either strain *A* or strain *B* (or both). Examples of such situations are shown in Figure 1. In the absence of mutation (*µ*_*A*_ = *µ*_*B*_ = 0) one has Φ_*A*_ *=* Φ_*B*_ *=* 0, and 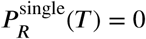 trivially as mutants never appear.

**Figure 1:**
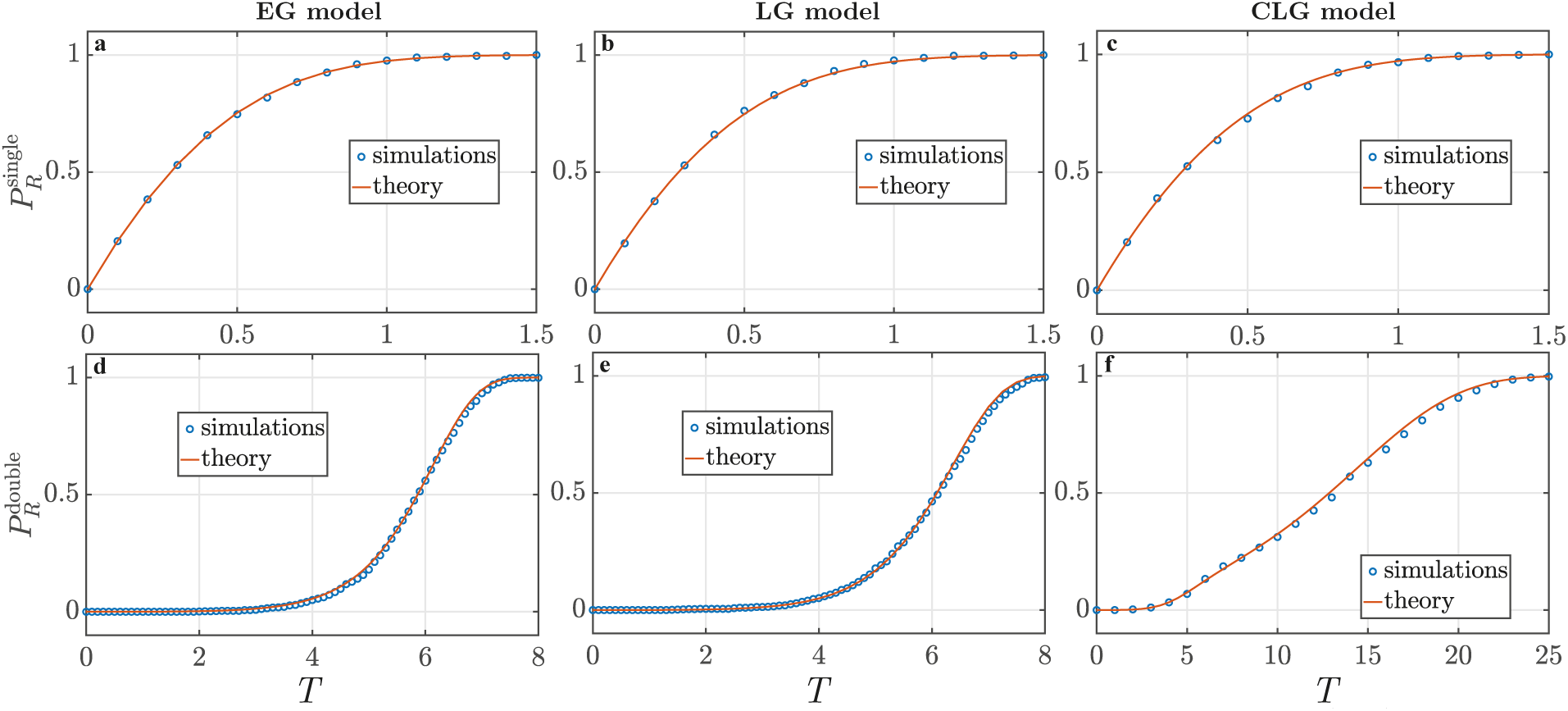
Single and double resistance probabilities for the three different growth models (EG exponential growth, LG logistic growth without competition between strains, CLG logistic growth with competition between strains). Theoretical predictions (solid lines) were obtained from equations (13) and (16), while numerical simulations (circles) were conducted using the Gillespie algorithm, results are averaged over 10000 runs. Parameters used: *b*_*S*_ = 1.1, *b*_*A*_ = 1.2, *b*_*B*_ = 1.3, *b*_*D*_ = 1.4, *d*_*S*_ = *d*_*A*_ = *d*_*B*_ = *d*_*D*_ = 0.1, *µ*_*A*_ = *µ*_*B*_ = 10^−4^, *n*_0_ = 10^4^. For the logistic growth models, we have set *ρ*_*i*_ = *b*_*i*_ − *d*_*i*_ for *i* ∈ {*S, A, B, D*}, and *k*_*S*_ = 10^6^, *k*_*A*_ = 1.1 × 10^6^, *k*_*B*_ = 1.2 × 10^6^, *k*_*D*_ = 1.3 × 10^6^.

### 3.3. Double resistance

The procedure to obtain the probability of double resistance is analogous to that for single resistance. The main difference is now that there are two sources, strains *A* and *B*. The probability of having at least one double-resistant cell at time *T* becomes (see Section S1.2.2 of the Supplementary Material) 

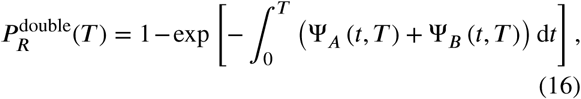

with 

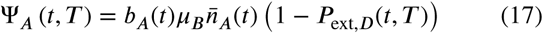

and 

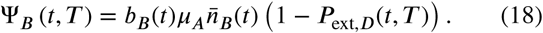

The terms Ψ_*A*_(*t, T*) and Ψ_*B*_(*t, T*) represent the rates with which double-resistants are produced by mutation of strains *A* and *B* at time *t*, respectively, and requiring that their lineage survives until time *T*. We will refer to 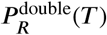 in Eq. (16) as the *probability of double resistance*.

We stress again that we are using the mean numbers of source cells as an input for these expressions (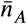 and 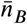). The limitations of this approach are discussed in Section 6.

### 3.4. Numerical integration of the probabilities of resistance

The previous expressions for the probabilities of resistance cannot always be reduced further. In addition, if the birth or death rates depend on the cell number of one or more strains, it is necessary to make a deterministic approximation (involving the replacement of the rates by their mean values) in order to compute the integrals in Eqs. (11), (13), and (16). In many cases it is not possible to do this analytically, and one has to resort to evaluating the integrals numerically.

Evaluating the probabilities of single or double resistance at time *T* requires the calculation of the functions Φ_*A*_(*t, T*), Φ_*B*_(*t, T*) and Ψ_*A*_(*t, T*) and Ψ_*B*_(*t, T*) in Eqs. (13) and (16) for all times *t* up to *T*. These objects in turn involve *P*_ext,*i*_(*t, T*) for *i* ∈ {*A, B, D*}, and evaluating these would require access to the objects *β*_*i*_(*t, t*′) for all combinations of times *t, t*′ up to *T* [Eq. (11)]. We recall that 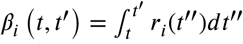 is the integrated net growth rate for strain *i*, see Eq. (12). As a consequence we would have to integrate the net growth rate *r*_*i*_ over the intervals [*t*′, *t*] for all combinations of *t*′ < *t* in the range up to *T*.

To make this process more efficient we first notice that – in absence of additional production of strain *i* through mutation – one has 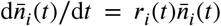. As a consequence 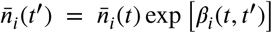. If we set 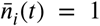, then the quantity *β*_*i*_(*t, t*′) can directly be expressed in terms of 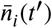, 

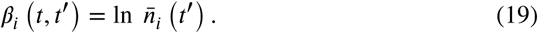

Therefore, the calculation of *β*_*i*_(*t, t*′) for a fixed combination of *t* and *t*′ (*t*′ > *t*) reduces to the problem of finding 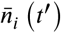 for the initial condition 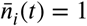. In principle this can be obtained from numerically integrating the growth law for strain *i* from the deterministic equations for 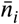, but it would imply separate integration runs for each initial time *t* due to the required initial condition 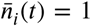. For the model with competition between strains, this also involves integrating the growth laws for all other strains.

In order to streamline this approach, our strategy consists of expressing the quantity 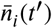 (with initial condition 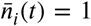) in terms of the solution 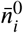 of the deterministic mean growth law for strain *i* with initial condition 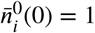 at time 0. The details of this are described in Section S3.2 of the Supplementary Material.

For the LG model we find (*t*′ > *t*), 

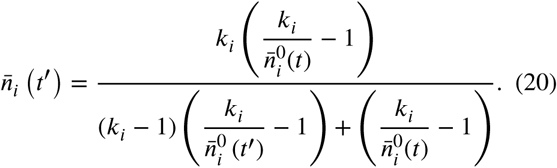

We re-iterate that this is the solution of the growth law for strain *i* for times *t*′ > *t*, subject to the initial condition 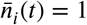. This initial condition can be verified directly from Eq. (20). We stress that 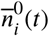 and 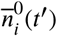 both feature on the right-hand side of Eq. (20).

Combining Eq. (20) with Eq. (19) we can find the integrated growth rate *β*_*i*_(*t, t*′)for all combinations *t* < *t*′ in the range up to *T*, provided we know the solution 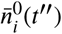 of the growth law for strain *i* with initial condition 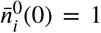 (*t*′′ < *T*). This trajectory can be obtained from one single numerical integration, significantly reducing the computational effort.

A similar approach can be taken for the model with competition between strains. This is discussed in more detail in Section S4 of the Supplementary Material. In the next section we proceed to analyse the probabilities of single and double resistance for each growth model and compare the theoretical predictions with results from numerical simulations. The simulations are carried out using the Gillespie algorithm (Gillespie (1977, 1976)). For processes with constant reaction rates this algorithm generates a statistically faithful ensemble of sample paths. For processes with time-dependent rates, the algorithm can still be used as an approximation. To do this we disregard the time dependence of the reaction rates during each Gillespie step, but then update the rates once a birth or death event has been executed. This is not an exact procedure, but the approximation works well if the rates do not change significantly during each Gillespie step. For a more detailed discussion of the algorithm and its limitations see Masuda and Rocha (2018) and Anderson (2007).

We focus on studying the probabilities of single and double resistance for the three growth models, and we study cases with constant drug and time dependent drug concentrations, respectively. We will concentrate on the effects the choice of the growth model has on these probabilities.

## 4. Probability of resistance for growth with constant coefficients

### 4.1. Setup, and comparison of theory and simulation

In the case of constant drug concentrations (approximating e.g. constant intravenous infusion, Shargel et al. (2004)) the birth and death rates for the exponential growth model [*b*_*i*_ and *d*_*i*_ in Eq. (10)] are time-independent. Similarly, the quantity *ρ*_*i*_ in the logistic growth models does not explicitly depend on time. Any dependence of *b*_*i*_ and *d*_*i*_ on time in the logistic models is through a functional dependence on 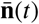 [c.f. Eqs. (7) and (8)]. To establish the baseline behaviour of the different growth models, we address this case of constant coefficients *b*_*i*_ and *d*_*i*_ in this section, that is coefficients without explicit dependence on time. Specifically, we will study how the probabilities of single and double resistance depend on the key model parameters for the three different types of growth.

The theoretical predictions of single and double resistance are obtained from Eqs. (11), (13) and (16). The extinction probability, the probability for single and double resistance can always be expressed in terms of the trajectories 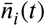, although the remaining integrals in the above equations may have to be carried out numerically.

For constant coefficients, closed-form solutions can be obtained for some of the quantities we are interested in, and in other cases we proceed numerically: (i) For the EG model, we can find a closed-form solution for the extinction probabilities *P*_ext,*i*_(*t, T*), this is given in Eq. (S15) in the Supplementary Material. Additionally, the mean cell numbers 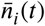 for each strain can be obtained in closed form from Eqs. (1), see Eq. (S14). With these solutions in turn, one can then derive closed-form expressions for the functions Φ_*A*_, Φ_*B*_, Ψ_*A*_ and Ψ_*B*_, and the probabilities of single and double resistance. This is explained further in Section S2 of the Supplementary Material. (ii) For the LG model, we can only find a closed-form solution for 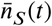, but not for the other strains. The probabilities of single and double resistance are calculated numerically using Eqs. (13) and (16). For further details see also Section S3 of the Supplementary Material. (iii) For the CLG model, we cannot express the mean cell number in closed form for any of the strains. This is due to the coupling and non-linearity of the equations for the mean cell numbers. The probabilities of single and double resistance are again obtained numerically.

For the time-dependent dosing protocols considered in Section 5, we have not been able to find closed-form solutions of Eqs. (1) for any of the strains and in any of the growth models. As a consequence the extinction probabilities and probabilities of single or double resistance cannot be found in closed form either, and have to be evaluated numerically.

As mentioned above in Section 2.4, in order to be able to compare the outcome of the different growth models, we specify fixed death rates *d*_*i*_ for each strain, and then use these for all three types of growth. In the EG model also specify the birth rates *b*_*i*_. In the two logistic growth models we then set *ρ*_*i*_ ≡ *b*_*i*_ − *d*_*i*_, so that the behaviour of all three models is similar for very small cell numbers.

Figure 1 illustrates a typical profile of the probabilities of having at least one single-resistant individual (either of type *A* or *B*) or at least one double-resistant individual, respectively. In the figure we compare the theoretical predictions for the three growth models against results from numerical simulations of the stochastic dynamics. The parameters used in the figure are for illustration, and do not necessarily represent a realistic situation. They describe a scenario in which the net birth-death rates, *r*_*i*_ = *b*_*i*_ − *d*_*i*_, are positive for all strains. This ensures that cell numbers increase in time (on average), so that single and double mutants eventually emerge. As the graphs show, the profile of the emergence of single resistance is very similar for the three different growth models, while double resistance tends to emerge later in the model with competition than in the other two scenarios. Further illustrations comparing simulation and theory for different choices of the model parameters are shown in the Supplementary Material (Section S5.2).

In the following sections we discuss the behaviour of both single and double resistance when varying different model parameters. In particular, we consider the outcome as a function of the carrying capacities, *k*_*i*_. In the limit of very high carrying capacities (very abundant resources), the logistic models reduce to the exponential model with unlimited growth. When the carrying capacities are finite, growth becomes restricted due to limited availability of resources.

### 4.2. Single resistance

Single resistance emerges through mutations during reproduction events of the sensitive strain *S* and is subsequently maintained provided the mutant strain does not become extinct. When the carrying capacity of sensitive cells *k*_*S*_ and single-resistant cells *k*_*A*_ and *k*_*B*_ are high compared to the initial number *n*_0_ of sensitive cells, there is no noticeable difference between the predictions of single resistance in the three different growth models (all three models lead to largely exponential growth); an example is shown in the upper row of Figure 1. However, as we will discuss below, the relation between *k*_*S*_ on the one hand, and *k*_*A*_ and *k*_*B*_ on the other, can affect the timing of the emergence of single-resistants in the competitive model. This is the case when these coefficients are close to each other, and to the the initial number of cells.

We show theoretical predictions for single resistance in Figure 2. In the figure we focus on the model with exponential growth, and we show the probability of finding at least one resistant cell as a function of time *T*, and varying a selection of model parameters. High probabilities of resistance are indicated by light colours. As one would expect, mutants tend to appear sooner as the birth rate *b*_*S*_, the initial number of sensitive cells *n*_0_ or the mutation rates *µ*_*A*_, *µ*_*B*_ are increased. This can be seen by the increased amount of lighter colours as one moves up along the vertical axes of panels (a), (c) and (d) in Figure 2. Panel (b) shows that the probability of single resistances decreases for increasing death rates *d*_*S*_ of the sensitive strain. Figure 3 illustrates how the predictions for single resistance differs between the logistic models with and without competition between strains (LG and CLG respectively), and when the carrying capacity *k*_*S*_ is varied. In the LG model single mutants tend to appear sooner when the carrying capacity *k*_*S*_ is high, see Figure 3 (a). This is as expected, as for higher values of *k*_*S*_ the limitations of growth of the sensitive strain only set in at large cell numbers. The resulting higher number of sensitive individuals increases the chance of producing single mutants.

**Figure 2:**
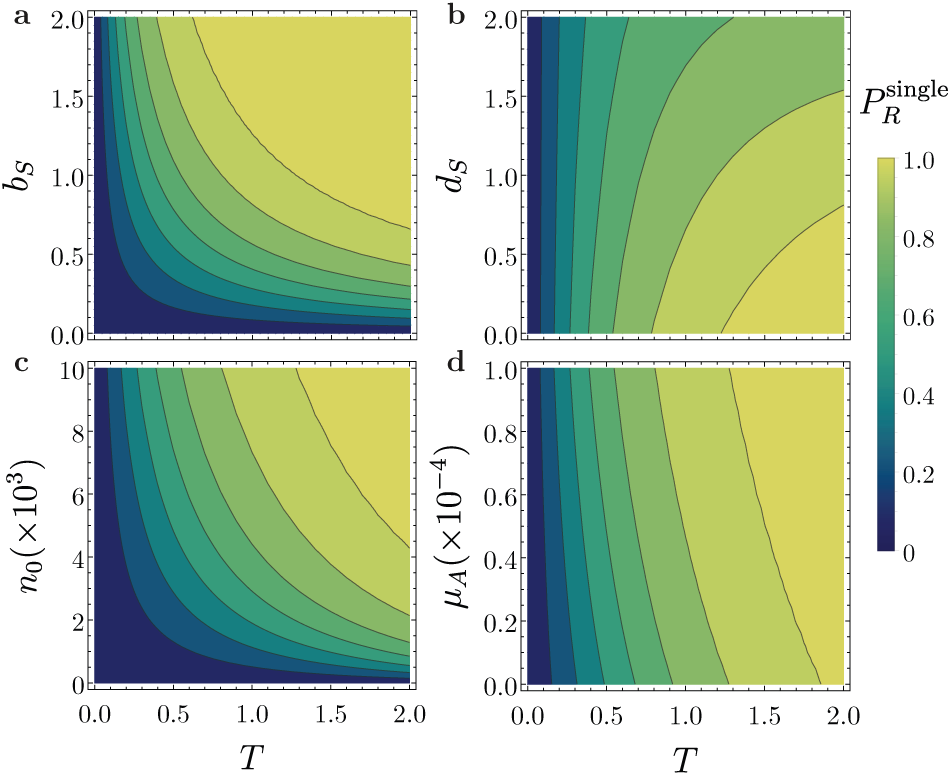
Theoretical prediction of the probability of single resistance obtained from Eq. (13) for the exponential growth model when varying only one parameter while keeping the others constant. When not varied, the parameters used are *b*_*S*_ = 1.0, *b*_*A*_ = 1.1, *b*_*B*_ = 1.2, *b*_*D*_ = 1.3, *d*_*S*_ = *d*_*A*_ = *d*_*B*_ = *d*_*D*_ = 0.1, *µ*_*A*_ = *µ*_*B*_ = 10^−4^, and *n*_0_ = 10^4^.

**Figure 3:**
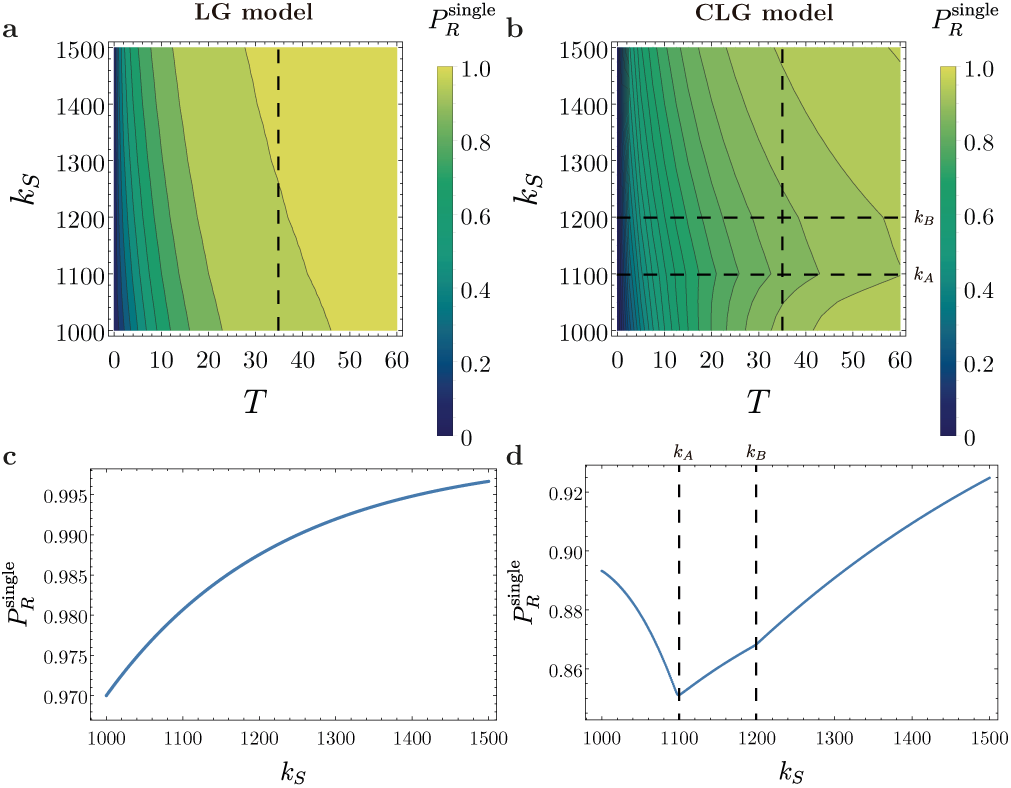
Theoretical prediction of the probability of single resistance for the logistic growth models without competition [LG, panels (a) and (c)], and with competition [CLG, panels (b) and (d)]. This is shown as a function of the carrying capacity *k*, for parameters *ρ*_*S*_ = 0.9, *ρ*_*A*_ = 1.0, *ρ*_*B*_ = 1.1, *ρ*_*D*_ = 1.2, *d*_*S*_ = *d*_*A*_ = *d*_*B*_ = *d*_*D*_ = 0 1, *µ*_*A*_ *=* 10^−3^, *µ*_*B*_ *=* 10^−4^, *n*_0_ = 10^3,^ *K*_*A*_ *=* 1.1 × 10^3,^ *K*_*B*_ *=* 1.2 × 10^3,^ *K*_*D*_ *=* 1.3 × 10^3^ Panels (c) and (d) show vertical cuts in panels (a) and (b) respectively, indicated by the vertical dashed line in the upper graphs at *T* = 35.

The dependence of the probability of single resistance on *k*_*S*_ is more intricate in the model with competition between strains (CLG). If the carrying capacity for strain *S* is close to the initial number of sensitive cells in the population, then the probability that single mutants emerge depends on how *k*_*S*_ compares to the carrying capacities *k*_*A*_ and *k*_*B*_ of the single mutant strain. We illustrate this Figure 3 (b) and (d) for a case with *µ*_*A*_ > *µ*_*B*_ and *k*_*B*_ > *k*_*A*_. The former of these conditions implies that the first single mutants are likely to be of type *A*.

The curve in Figure 3 (d) shows interesting behaviour: We find that the probability of single resistance decreases with increasing *k*_*S*_, provided *k*_*S*_ is sufficiently small. At an intermediate value of *k*_*S*_ ≈ *k*_*A*_ we observe a minimum. For higher values of *k*_*S*_, the probability of single resistance then increases again with *k*_*S*_. A small ‘kink’ is seen at *k*_*S*_ ≈ *k*_*B*_.

We attribute these features to a combination of several counteracting effects: (i) Generally, an increase in the carrying capacity *k*_*S*_ leads to a less restricted and therefore faster growth of the sensitive strain. A high carrying capacity also leads to higher numbers of sensitive cells in the long run. Both of these effects favour the emergence of single-resistant cells. (ii) If the carrying capacity *k*_*S*_ is lower than *k*_*A*_ and *k*_*B*_, then the total cell number *n*_*T*_ is lower than *k*_*A*_ and *k*_*B*_ when strain *S* saturates. The population consists mostly of sensitive cells at this point. As first mutants emerge, they will be able to proliferate until the total size of the population reaches the carrying capacity of the relevant single-mutant strain. This is likely to be *A* because of the higher mutation rate *µ*_*A*_ > *µ*_*B*_. The maximum number of single mutants *A* that can arise is broadly governed by the difference between *k*_*A*_ and the number of sensitive cells. The latter in turn is approximately given by *k*_*S*_. Therefore, as *k*_*S*_ approaches *k*_*A*_ from below there is only little room for single-resistant cells to proliferate before saturation sets in. In some cases only a few single-resistant cells can be produced, and these may eventually die due to stochastic fluctuations. This effect acts to reduce the probability of single resistance as *k*_*S*_ approaches *k*_*A*_ from below.

In the example of Figure 3 (d), the second effect dominates over the first in the regime *k*_*S*_ < *k*_*A*_, and the probability of single resistance is a decreasing function of *k*_*S*_. When *k*_*A*_ < *k*_*S*_ < *k*_*B*_, the suppression of growth of mutant strain *A* (effect (ii) above) is no longer relevant. Effect (i) now dominates, and the probability of single resistance becomes an increasing function of *k*_*S*_. As a result a minimum is found for the probability of single resistance at *k*_*S*_ = *k*_*A*_. When *k*_*S*_ hits *k*_*B*_ any suppression effect due to mutants of type *B* is also removed, and the probability of single resistance increases more sharply with *k*_*S*_. This results in a ‘kink’ at *k*_*S*_ = *k*_*B*_ in Figure 3 (d).

### 4.3. Double resistance

We now compare the probability of double resistance in the three growth models. In Figure 4 we show the effects of varying *b*_*S*_, *d*_*S*_, *b*_*A*_, and *n*_0_ in turn; for other parameters see Section S5.1 of the Supplementary Material. In general, the logistic growth model without competition between strains (LG) seems to produce largely similar behaviour as the model with exponential growth (compare panels (a)–(d) with (e)–(h)). However, differences between the EG and LG models become apparent where the carrying capacities are varied. This will be discussed below in Fig. 5.

**Figure 4:**
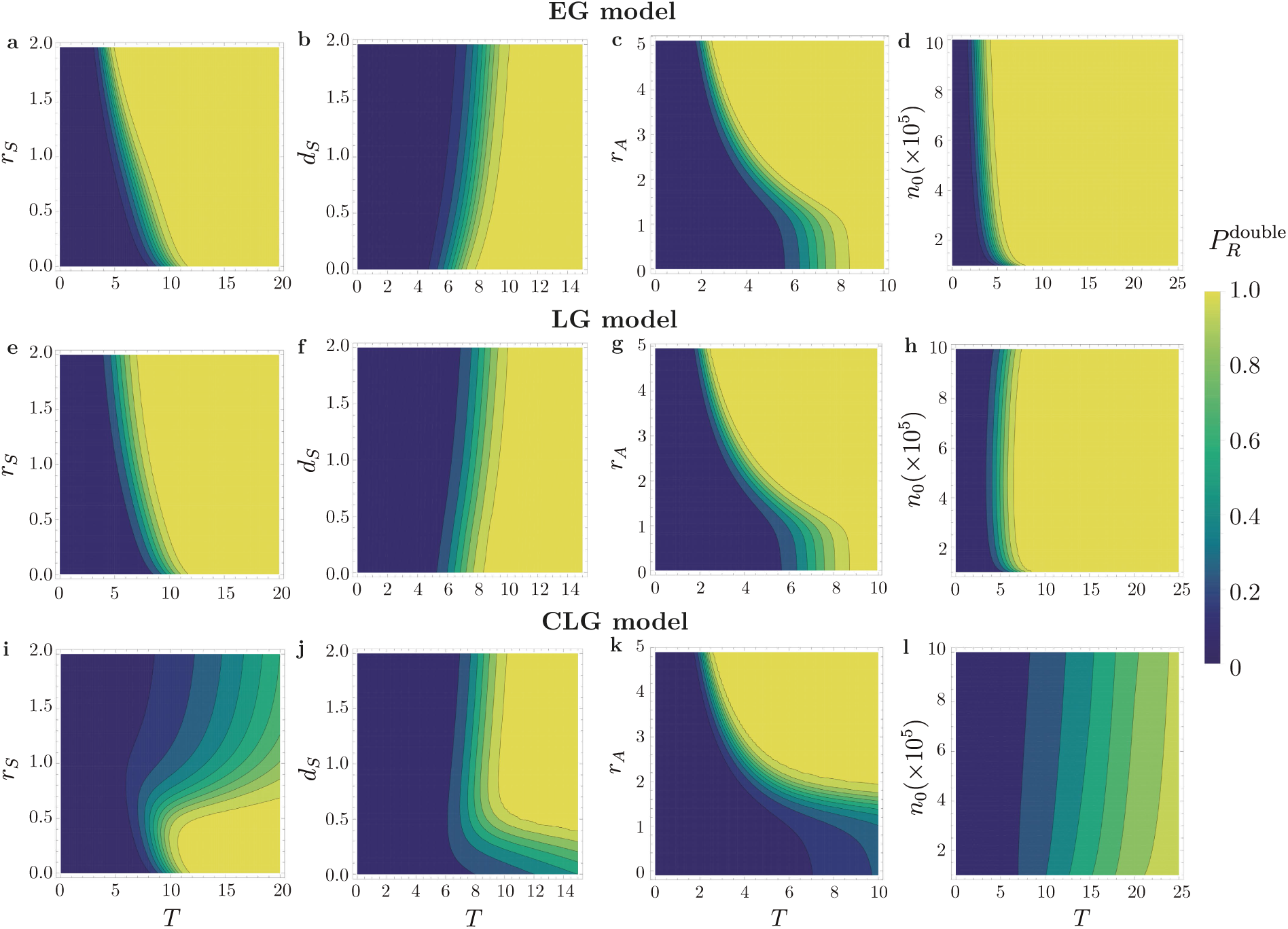
Theoretical prediction of the probability of double resistance for the three growth models when varying one parameter (EG exponential growth, LG logistic growth without competition between strains, CLG logistic growth with competition between strains). When not varied, the parameters used are *b*_*S*_ = 1.0, *b*_*A*_ = 1.1, *b*_*B*_ = 1.2, *b*_*D*_ = 1.3, *d*_*S*_ = *d*_*A*_ = *d*_*B*_ = *d*_*D*_ = 0.1, *µ*_*A*_ = *µ*_*B*_ = 10^−4^, and *n*_0_ = 10^4^. When varying *r*_*S*_ = *b*_*S*_ − *d*_*S*_(*r*_*A*_ = *b*_*A*_ − *d*_*A*_), only *b*_*S*_(*b*_*A*_) changes, keeping *d*_*S*_(*d*_*A*_) constant. As described in the main text, we have used *ρ*_*i*_ = *r*_*i*_ for the logistic growth models. Carrying capacities used: *k*_*S*_ = 10^6^, *k*_*A*_ = 1.1 × 10^6^, *k*_*B*_ = 1.2 × 10^6^, *k*_*D*_ = 1.3 × 10^6^. The lightest colour represents values of 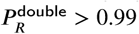.

**Figure 5:**
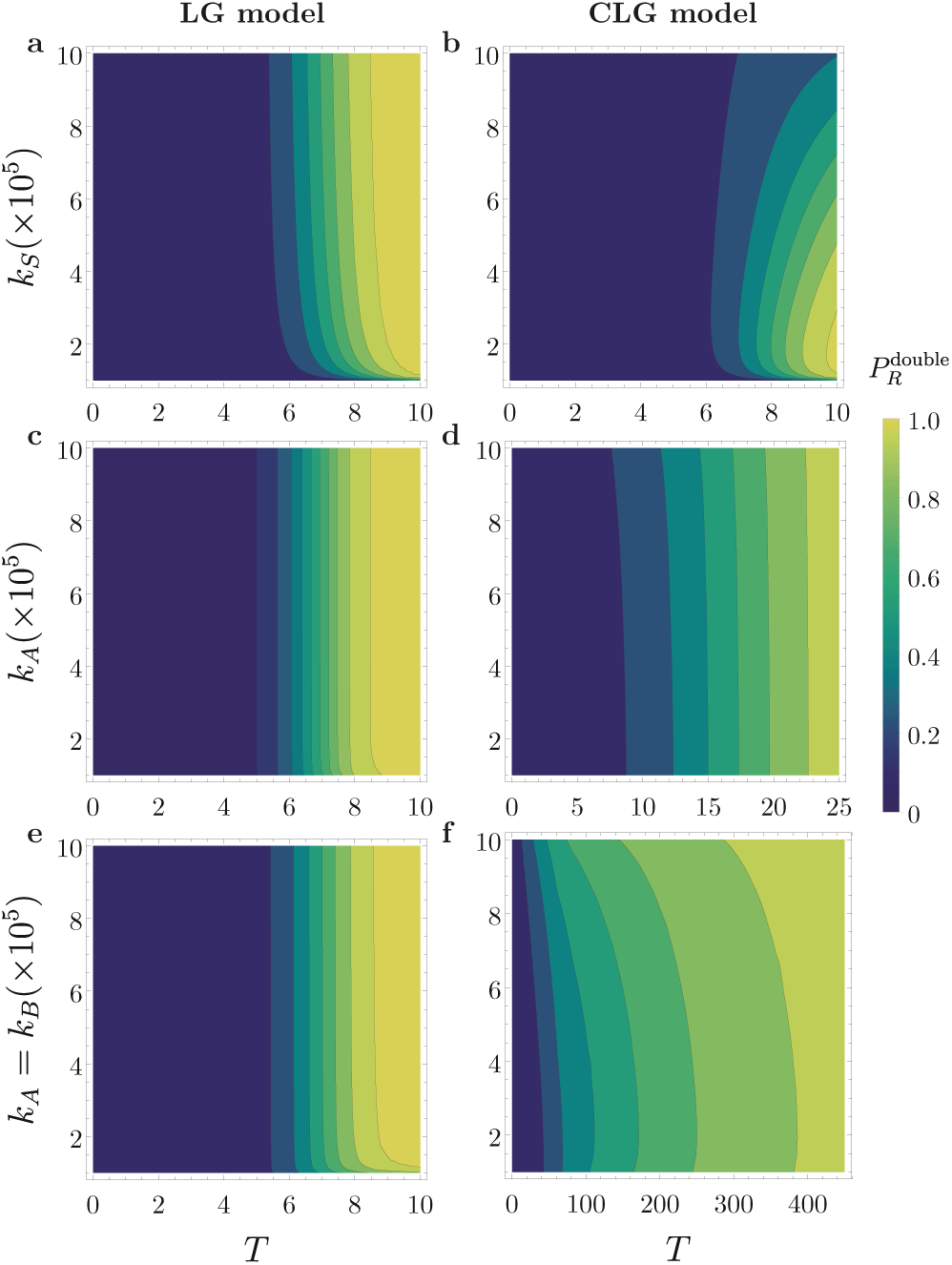
Theoretical prediction for the probability of double resistance for the logistic models without competition between strains (LG), and with between-strain competition (CLG) when varying the carrying capacities. When not varied, the parameters used are the same as in Figure 4. In panels (e) and (f), both *k*_*A*_ and *k*_*B*_ are varied at the same time. Notice the different range of times in panels (d) and (f) compared to the other panels.

The logistic model with competition between strains, however, shows notably different behaviour from the other two models [see Figure 4(i)-(l)]. In particular we make the following observations:

1. The time it takes for the probability of double resistance to reach a specified value near one in the CLG model is a non-monotonous function of the net growth rate *r*_*S*_. This can be seen from the curved shape of the contour lines in Figure 4 (i). It becomes more likely to observe double resistance at a given time for increasing *r*_*S*_, provided *r*_*S*_ is sufficiently small. Once *r*_*S*_ has reached a certain point, however, increasing *r*_*S*_ further reduces the probability of finding double resistance at a fixed time. We first look at the regime of small *r*_*S*_. The total cell number remains low enough to avoid saturating the growth of the emerging single mutants. Double-resistants then tend to appear sooner the larger the growth rate *r*_*S*_ of the sensitive strain. Hence, the probability of double resistance increases with *r*_*S*_. As *r*_*S*_ is increased further, strain *S* grows fast enough for the total cell number to saturate the proliferation of emerging single-resistant strains (due to the competition between strains). This makes the emergence of double mutants less likely, hence the probability of double resistance decreases with *r*_*S*_.
2. For low values of the death rate of strain *S, d*_*S*_, double mutants tend to appear later in the CLG model than in the other two growth models [compare the lower right of panel (j) in Figure 4 with the data in the lower right of panels (b) and (f)]. This effect is similar what was discussed in item 1. above. Lowering the death rate *d*_*S*_ while keeping the birth *b*_*S*_ fixed results in faster net growth of the sensitive strain. This means that the growth of the single-resistant strain saturates, thus reducing the chance of emergence of double mutants.
3. For low values of the net growth rate *r*_*A*_ of strain *A* (accounting for birth and death), double mutants tend to appear very late in the CLG model compared to the other growth models [compare the lower right of Figure 4 (k) with those in panels (c) and (g)]. A low value of *r*_*A*_ means that more time is required to generate a number of *A* individuals which is sufficiently large to make the appearance of double mutants likely. As the other present strains (*S* and *B*) have more time to grow, the total population size *n*_*T*_ becomes large enough to saturate the growth of strains *A* and *B* before the first double-resistants emerge. As a consequence, the chance of double mutants to emerge decreases.
4. For an increased number of initial cells *n*_0_, it becomes less likely to observe double mutants at a given time [see Figure 4 (l)]. The delay in the emergence of double mutants is again explained by a saturation of the growth of sensitive and single-resistant strains. For higher initial number of sensitive cells, saturation occurs sooner, and double-resistant cells appear later.

These observations suggest that competition can affect the emergence of double resistance. The predictions shown in Figure 4 indicate that the typical time at which the first double-resistants appear can be notably different when competition is taken into account [compare e.g. Figure 4 panel (l) with panels (d) and (h)]. The difference between the competitive model and the other two models in producing double-resistants is mainly due the saturation of the sensitive and single-resistants before double mutants appear. This is an important factor to take into account for modelling resistance, in particular in situations where resources are limited. Moreover, our analysis indicates that the optimum treatment, i.e. the one that most delays the emergence of double-resistants, can also differ across the models. This can be illustrated using the behaviour of the probability of double resistance as *d*_*S*_ is varied, see Figure 4 (b), (f) and (j). The model with competition between strains predicts that a low *d*_*S*_ would maximise the delay the emergence of double mutants, while the other models predict that a higher *d*_*S*_ would be best.

In addition, the carrying capacities can also affect the probability of double resistance in both logistic growth models. An example of this is displayed in Figure 5, where we vary the carrying capacity of strain *S*, or of only one of the single-resistant strains (strain *A*), or of both (strain *A* and *B*) at the same time.

The lower end of panels (a) and (b) show a situation in which the coefficient *k*_*S*_ is above the initial number of cells *n*_0_ = 10000, but close to it. Double mutants tend to appear only after a relatively long time in this situation in both logistic models. When *k*_*S*_ is sufficiently far above *n*_0_, its precise value does not have a pronounced effect on double resistance in the LG model [see Figure 5(a)]. In the CLG model, however, an increase in the carrying capacity *k*_*S*_ of the sensitive strain results in a delayed emergence of double mutants [Figure 5(b)].

When varying only *k*_*A*_ [Figure 5 (c) and (d)], the probability of double resistance does not show a notable change in either of the logistic models. Independently of the dynamics of strain *A*, double mutants can still be produced from strain *B*. Notice, however, that double mutants tend to appear at later times in the CLG model than in the LG model. When both carrying capacities *k*_*A*_ and *k*_*B*_ are varied together [panels (e) and (f)], double resistance is significantly delayed in the CLG model in comparison to the LG model when *k*_*A*_ and *k*_*B*_ are high.

In summary we conclude that the carrying capacity of strain *S* has a stronger effect on the emergence of double resistance than each of the single-resistant carrying capacities.

## 5. Time-dependent drug concentrations

The analysis in the previous section focused on situations in which model parameters do not vary in time. In many instances however, drugs are administered using time-varying dosing schedules, giving rise to time-dependent death and growth rates. Most drug therapies use periodic dosing schedules, resulting in periodic time-dependences of drug concentrations (see e.g. Ibrahim et al. (2004); Bur-nette (1992)). Modelling the the pharmacokinetics of different methods of repeat drug administration, we consider two different types of time-dependence for the drug concentrations. In the first scenario, drug concentrations are continuous in time and follow a sinusoidal profile. This approximates the pharmacokinetics for example of repeat antibiotic dosing via an extravascular route (e.g. oral administration). In the second scenario the drug concentration profiles consist of a periodic series of pulses, representing, for example, intermittent intravenous boluses (Shargel et al., 2004; Chambers, 2019).

We note that the net growth rates of the strains affected by the drugs (*r*_*i*_ = *b*_*i*_ − *d*_*i*_ for *i* ∈ {*A, B, D*}) in Eq. (10) can become negative when the concentration of the relevant drugs are sufficiently high. This leads to an effective reduction of the number of cells of the affected strains. Motivated by clinical treatment protocols, the drug therapies discussed below are designed such that the sensitive strain will eventually become extinct, which is the goal of treatment [see e.g. Foo and Michor (2009)]. This is achieved if 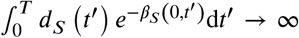 as *T* → ∞, with 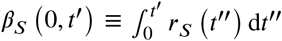 as explained in Section 3.1. To achieve this for oscillatory drug concentrations with period *P* it is sufficient that *β*_*S*_ (0, *P*) < 0, i.e., that the time-average of the net growth rate over one period is negative (assuming that *d*(*t*′) remains above a non-zero value for all *t*′).

### 5.1. Sinusoidal drug concentrations

We first consider a sinusoidal drug profile, with period *P*. The drug concentrations take the form 

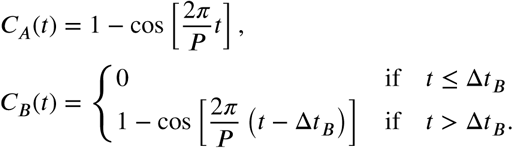

Drug 𝒜 is applied starting from time *t* = 0, initially with concentration zero (*C*_*A*_(*t* = 0) = 0), and then following a periodic profile. The application of drug ℬ starts at a later time Δ*t*_*B*_(*C*_*B*_(*t*) = 0 for *t* ≤ Δ*t*_*B*_). The concentration of ℬ then also follows a sinusoidal profile from time Δ*t*_*B*_ onwards. The factors *f*_*A*_ and *f*_*B*_ range from 1 to 3 over each cycle. The drug profiles *C*_*A*_ and *C*_*B*_ are shifted by a time Δ*t*_*B*_ [Δ*t*_*B*_ mod *P* when Δ*t*_*B*_ > *P*] and are out of phase by a fraction Δ*t*_*B*_/*P* of a period [(Δ*t*_*B*_ mod *P)* /*P* when Δ*t*_*B*_ > *P*].

A sample profile of the drug concentrations is shown in Figure 6 (a) for *P* = 1 and Δ*t*_*B*_ = 0:5; the corresponding growth rates for the different strains are shown in Figure 6 (c).

**Figure 6:**
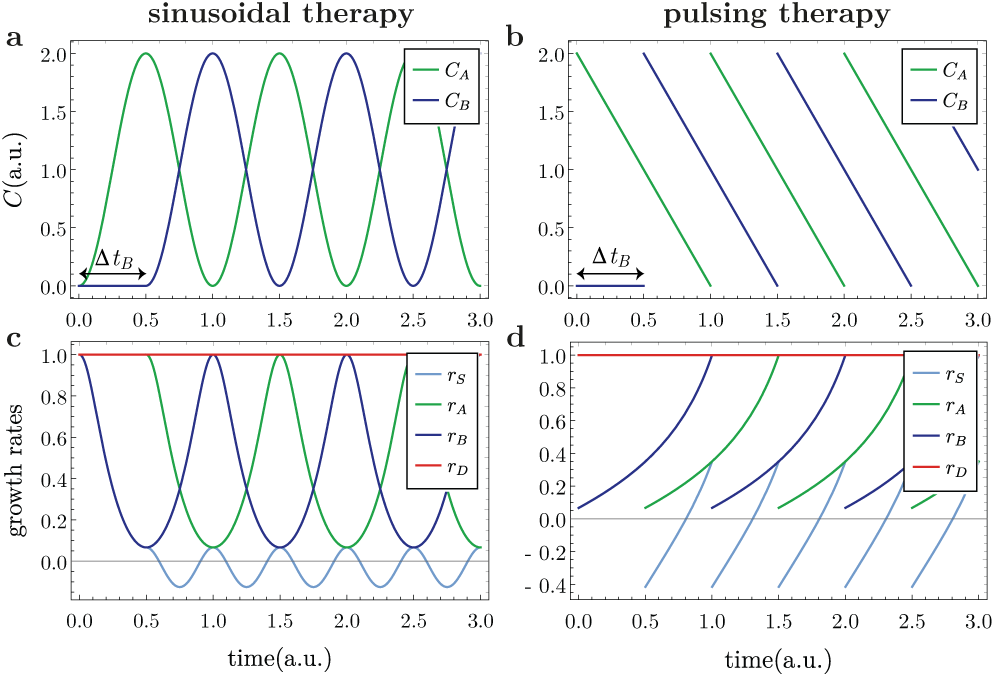
Drug concentrations *C*_*A*_(*t*) and *C*_*B*_(*t*) in the sinusoidal protocal (a) and for pulsing profiles (b) (period *P* = 1 and Δ*t*_*B*_ = 0.5). Panels (c) and (d) show the respective growth rates (*r*_*i*_(*t*) = *b*_*i*_(*t*) − *d*_*i*_(*t*), with *i* ∈ {*S, A, B, D*}) affected by the drug effect described in equations (9)-(10). Parameters used: 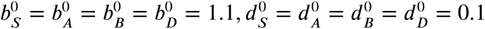.

### 5.2. Pulsing drug concentrations

In this protocol, the concentration 𝒜 of drug is assumed to be of the form 

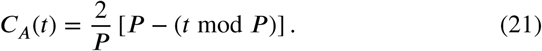

This means that the profile *C*_*A*_(*t*) starts off at *C*_*A*_(*t* = 0) = 2, and then falls linearly with slope 2*P* ^−1^, so that the drug concentration becomes zero at *t* = *P*. Then the next pulse starts, i.e., *C*_*A*_ is re-set to its maximum value, and then falls off linearly again. Similar profiles have been used in Ibrahim et al. (2004); Chakrabarti and Michor (2017).

The concentration of drug ℬ is assumed to be zero up to time Δ*t*_*B*_, and then follows a similar sequence of pulses. This is described by 

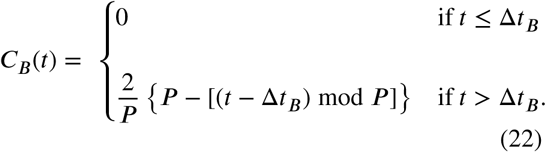

The concentrations of both drugs are out phase as in the sinusoidal drug therapy, and attain their minima at the same times as in the sinusoidal protocol. Figure 6 (b) illustrates a sample profile for *P* = 1 and Δ*t*_*B*_ = 0.5; we show the corresponding net growth rates for the different strains in panel (d).

### 5.3. Probability of double resistance for time-dependent dosing schedules

In Figure 7 we show the probability of double resistance resulting from the periodic dosing protocols. Data is presented as a function of time, and for any pairing of the two therapy protocols and the three different growth models. We have considered the cases where drug ℬ is administered from times Δ*t*_*B*_ = 0.0, 0.5 *P* and 1.0 *P*, with *P* = 1. Predictions for other values of Δ*t*_*B*_, as well as the probability of single resistance, can be found in Section S5.3 in the Supplementary Material. We have chosen parameters such that the carrying capacities are not too far from the initial number of sensitive cells, allowing the limitations on growth in the logistic models to set in quickly. We observe that the double-resistant strain tends to appear later in the pulsing therapy than in the sinusoidal therapy (to see this compare the value of 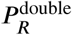 for the two protocols at a given time). This is not surprising as, for equal parameters, the growth rate *r*_*S*_ spends more time taking negative values in the pulsing therapy than in the sinusoidal therapy [compare panels Figure 6 (c) and (d)].

**Figure 7:**
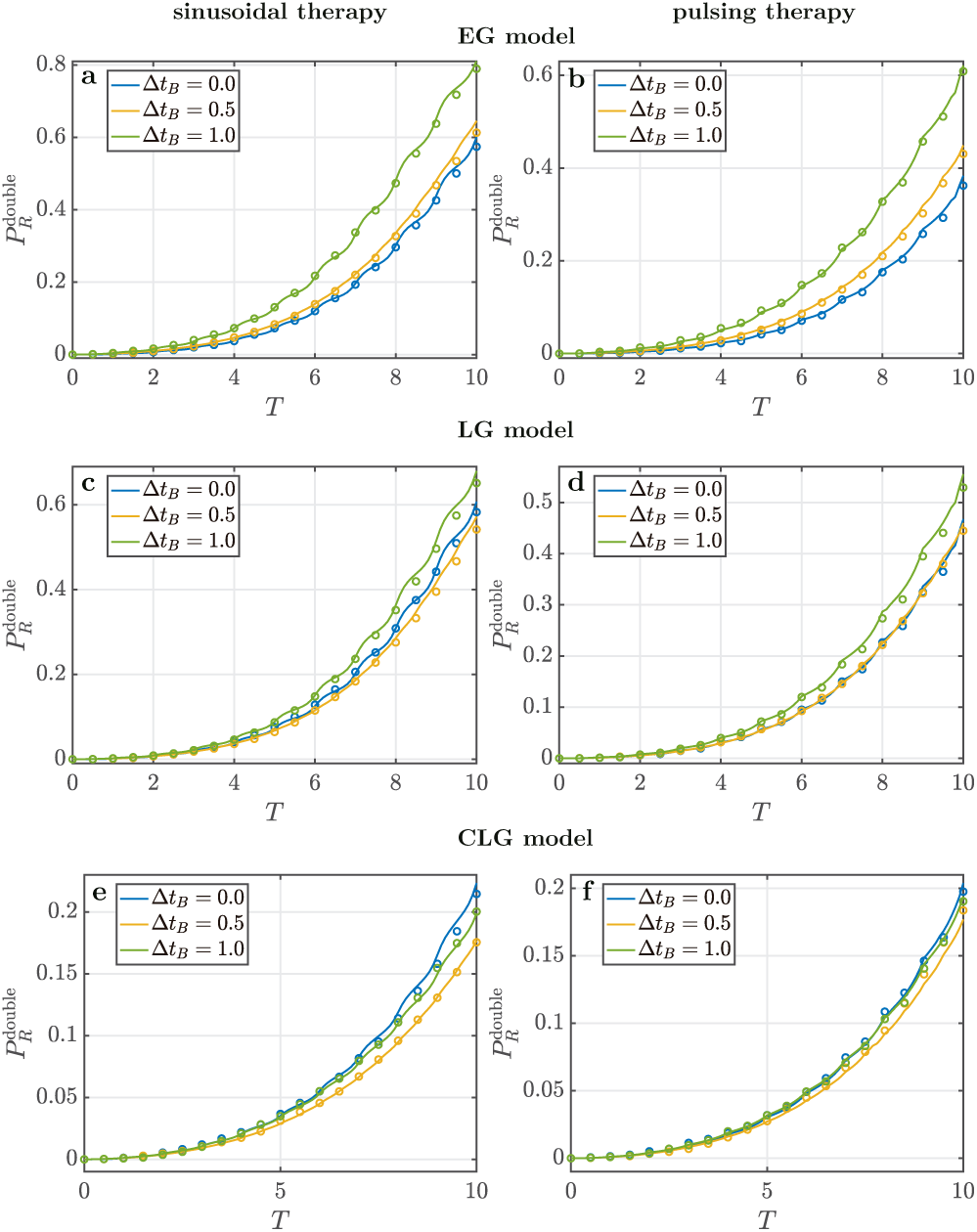
Probability of double resistance as function of time for each of the three growth models for different values of Δ*t*_*B*_ for sinusoidal and pulsing therapies (EG exponential growth, LG logistic growth without competition between strains, CLG logistic growth with competition between strains). Solid lines represent the theoretical predictions, while open circles the numerical simulations. Parameters used: 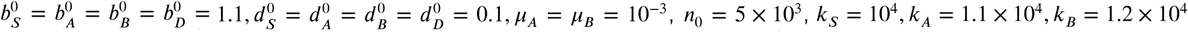, and *k*_*D*_ = 1.3 × 10^4^.

In addition, we also note that, at equal times after the treatment has started, the probability of double resistance is considerably lower in the logistic model with competition between strains than in the other two growth models, independently of the treatment protocol. This is due to the early saturation of the growth of the single-resistant strains in the CLG model, inhibiting the emergence of double-resistants. We observe that the fraction of sensitive cells in the total population is small at the point where the growth of single-resistants saturates due to the effects of drugs. This is because the growth of sensitive cells is heavily suppressed by the treatment. As a consequence, the contribution of sensitive cells to the saturation of single-resistant strains is negligible in the CLG model.

Interestingly, the data in Figure 7 also shows that the optimum treatment within our model setting, i.e., the treatment with the lowest probability of double resistance at a given time, can be different for different growth models. We describe this in the context of the sinusoidal therapy: the optimum treatment in the exponential growth model is obtained when both drugs are administered at the same time, i.e. when Δ*t*_*B*_ = 0 [see Figure 7 (a)]. This is not the case in the other two growth models [see panels (c) and (e)]. In fact, for the logistic model with competition between strains [panel (e)], the case Δ*t*_*B*_ = 0 turns out to be the worst treatment of the ones shown here, as double mutants would appear earliest. For the pulsing therapy, we observe similar behaviour. For both logistic growth models, the optimum treatment in each therapy is found when Δ*t*_*B*_ = 0.5, i.e., when the drugs are applied alternately. This result is consistent with the findings from Peña-Miller et al. (2012) for synergistic drug therapies.

In Figure S7 of the Supplementary Material we re-plot the data of Figure 7, but as function of Δ*t*_*B*_ for a fixed time *T*. This further illustrates that the optimum treatment is achieved for different choices of the time lag Δ*t*_*B*_ in the different growth models. Another example showing that the optimum treatment varies across the growth models is presented in Figure S8 in the Supplementary Material for a scenario with higher carrying capacities.

## 6. Limitations due to deterministic approximation

The method we have presented to estimate the probabilities of single and double resistance can be used for constant and time-dependent drug concentrations in exponential and logistic growth models. However, our approach relies on an approximation. While we treat the emergence of single and double mutants and the persistence or extinction of their lineages as stochastic processes, we disregard fluctuations in the rates with which mutants first appear. To illustrate this, we focus on mutants of type *A*. They are arise from mutations in reproduction events of sensitive cells *S*. The rate with which individuals of type *A* emerge therefore depends on the number of sensitive cells in the population. As a consequence, the production of *A* in mutation events is a stochastic process with two levels of randomness. First, the number of sensitive cells is a random quantity. Second, each of the sensitive cells can produce mutants of type *A* with a certain probability upon reproduction, and the subsequent birth-death dynamics of the single-mutant cells is also stochastic. Our approach consists of neglecting the first type of randomness for the purposes of studying mutants of type *A*. In the production rate of *A* we replace the actual stochastic number of sensitive cells by its mean (over realisations). We do however retain the second type of randomness, and treat the subsequent extinction or persistence of the mutants as a stochastic process. This is captured by the extinction probabilities *P*_ext,*i*_(*t, T*) in Eq. (11). In very much the same way we treat the number of *A* and *B* cells as deterministic for the purposes of calculating probability of double resistance, but retain the stochasticity of the production of double mutants, and the subsequent evolution of their lineages.

This approximation works well for the estimation of the probability of single resistance. Single-resistant cells are produced from sensitive cells in mutation events, and the number of sensitive cells is large from enough from the beginning to neglect fluctuations.

Double resistance on the other hand is produced from single-resistant cells. Single-resistant cells are not present in the population at the beginning, and their numbers will initially be small. One would therefore expect that fluctuations of the number of cells of single resistants *A* and *B* will be more pronounced than those of the number sensitive cells. This may make the predictions for double resistance inaccurate. In simulations we find that this tends to be the case mostly when the first double mutant appears at long times after the start of the dynamics. This can occur for example when the initial number of sensitive cells, the mutation rates, or the birth rates are small. We compare the predictions of our calculations against simulations for such cases in Section S6 of the Supplementary Material.

Simulations confirm that the number of the single-resistant cells can be subject to significant fluctuations under these circumstances. As a consequence, the deterministic approximation in the production rates of double mutants becomes inaccurate, and with it the approximation for the probability of double resistance in Eq. (16). Joint effects of the random occurrence of single-mutants, and their subsequent growth are a possible explanation for the fluctuations of the number of single-resistants. Typical cases in which

The prediction for the presence single mutants appears to be affected less by these effects. We attribute this to the fact that single mutants arise from the sensitive strain, which is present from the start and not generated by random mutation events.

We note that the method we develop can be extended to more complex situations, such as models with heterogeneous competition. The approach can also be generalised to combination therapies of more than two drugs, as well as to cases with more complex interactions between the drugs. It may also be interesting to account for heterogeneity of the mutation rate within the population, as this can affect resistance (Alexander et al. (2017)). Given that the limitations of our approximation can be characterised, we think that the method we have developed can be useful for these questions, and for other biological problems in which multiple mutations occur sequentially.

## 7. Conclusions

Models for estimating the probability of drug resistance have previously been largely based on exponential growth (some exceptions are Austin and Anderson (1999); Baker et al. (2016)). The use of exponential growth models implies an assumption of infinite resources and the absence of competition between strains. Here, we have shown that the choice of growth model can affect the probability that resistant cells emerge in when resources are limited and when there is potential competition between strains.

To do this, we have investigated the evolution of single and double resistance in a stochastic multi-strain model. Our analysis focuses on three different growth laws: exponential growth, logistic growth without competition between strains and logistic growth with between-strain competition. We have examined cases in which the model parameters are not explicitly time-dependent, as well as simple time-dependent drug therapies. We have analytically estimated the probability of having at least one single or double-resistant cell in the population, and we have verified these predictions in numerical simulations. Our calculations require the evaluation of a large number of integral terms in the expressions for the probability of single and double resistance. As a by-product of our work we have provided strategies to reduce the number of integral terms to be evaluated, allowing us to make theoretical predictions more efficiently.

Our results show that the choice of growth model makes a difference for the probability of double resistance, both for abundant and limited resources—a distinction not seen for single drug resistance. Specifically, competition can considerably delay double resistance, both for constant model parameters and for time-dependent dosing schedules. This delay in double resistance occurs for a range of parameters for the sensitive and single-resistant strains. Consequently, modelling resistance with exponential growth laws may make combination treatments appear less effective than they are in the presence of between-strain competition. Further, careful attention needs to be paid to competition when planning treatments with time-dependent concentrations. Our results predict that the optimal dosing schedule for a combination treatment changes depending on the growth law used. An accurate representation of growth is therefore critical for infections where resource competition is strong. This is particularly relevant to long-term or chronic infections and cancer (Wang et al., 2003; Graham, 2008; Mideo, 2009; Hibbing et al., 2010; Baishya and Wakeman, 2019). The exact way in which growth is modelled therefore requires careful consideration in the design of drug therapies.

## Acknowledgements

We acknowledge support through a Presidential Doctoral Scholarship (The University of Manchester) to EBC, a Wellcome Trust Institutional Strategic Support Fund Award (204796/Z/16/Z) to DRG and TG. DRG is supported by a UKRI Innovation/Rutherford Fund Fellowship (MR/R024936/1). TG acknowledges financial support from the Maria de Maeztu Program for Units of Excellence in R&D, Spain (MDM-2017-0711).

## Data archiving

Data, scripts and source code are available on GitHub: https://github.com/ernestoberriosc/probabilities-resistance

## Author contributions

All authors contributed to conceiving and designing the study. EBC carried out the mathematical analysis and numerical simulations. All authors contributed to analysing results and data, and to writing the paper.

