## Supplementary Material for "Competition delays multi-drug resistance evolution during combination therapy"

Ernesto Berríos-Caro, Danna R. Gifford, and Tobias Galla

#### S1 Derivation of the probabilities of single and double resistance for general birth and death rates

In this section, we will give further details of the calculation of the probability of resistance, i.e., the probability of having at least one single resistant or double resistant individual at a certain time. We do this for general birth and death rates, assuming that these rates are given by deterministic functions. In other words, the birth and death rates may depend on the composition of the population, but if they do then we ignore any stochasticity of the number of different cells in the population. The population starts from $n_0$ sensitive cells at time $t = 0$, and with no other cells.

The method we will use is based on [1], where the probability of having at least one single resistant was calculated for an exponential growth model. We extend this method to obtain the probability of double resistants for each of the three growth models defined in the main text.

##### S1.1 Extinction probability for single cell and its lineage

We first consider the probability that one single individual of type $i$ present at time $t$ goes extinct by time $T$, along with its lineage. For strain $i$, this extinction probability is given by (see e.g. [2, 3]).

$$P_{\mathrm{ext},i}(t,T) = \frac{\displaystyle\int_t^T d_i\left(t'\right) e^{-\beta_i\left(t,t'\right)} \mathrm{d}t'}{1 + \displaystyle\int_t^T d_i\left(t'\right) e^{-\beta_i\left(t,t'\right)} \mathrm{d}t'}, \tag{S1}$$

with

$$\beta_i\left(t,t'\right) \equiv \int_t^{t'} r_i\left(t''\right) \mathrm{d}t'', \tag{S2}$$

where $b_i(t)$and $d_i(t)$ are the birth and death rates for cells of type $i$, and $r_i(t) = b_i(t) - d_i(t)$. This is an exact result provided that the birth and death rates are both given by deterministic functions.

##### S1.2 Probability of no resistance

The probability of having no resistant individuals of a given type $i \in \{A, B, D\}$ at time $T$ is obtained by dividing the interval $[0, T]$ into a large number $N$ of sub-intervals, each of length $\Delta t = T/N \ll 1$. One then requires that any resistant of type $i$ which is produced in a given sub-interval $[t_k, t_k + \Delta t] \subseteq [0, T]$ ($t_k = (k-1)\Delta t$, $k \in \{1, 2, \ldots, N\}$) goes extinct by time $T$, along with its lineage.
For any sub-interval there are two possibilities:

1. No relevant mutations occur, i.e. no resistant cells of type $i$ are produced. The probability that this happens is

$$1 - \overline{W}_i(t_k)\Delta t, \tag{S3}$$

as $\overline{W}_i(t_k)\Delta t$ is the probability that a relevant reproduction-mutation event occurs in this interval. The overbar indicates that we use a deterministic approximation for the rate $W_i(t_k)$.

2. A relevant mutation occurs (i.e., a resistant cell of type $i$ is produced), but it goes extinct along with its lineage by time $T$. The probability for this event is

$$\overline{W}_i(t_k)\Delta t P_{\text{ext},i}(t_k, T), \tag{S4}$$

with $P_{\text{ext},i}(t_k, T)$ as defined above.

Next, we will describe how to use this to estimate the probability of single and double resistance.

##### S1.2.1 Single resistance

Following this idea, the probability that either no single resistant cell arises in $[t_k, t_k + \Delta t]$, or if one arises it goes extinct by time $T$ along with its lineage becomes

$$\begin{aligned} P_{0,t_k}^{\text{single}}(T) &= \left[1 - \left(\overline{W}_A(t_k) + \overline{W}_B(t_k)\right)\Delta t\right] + \overline{W}_A(t_k)\Delta t P_{\text{ext},A}(t_k,T) + \overline{W}_B(t_k)\Delta t P_{\text{ext},B}(t_k,T), \\ &= 1 - \Delta t \Big\{ W_A(t_k)\left[1 - P_{\text{ext},A}(t_k,T)\right] + W_B(t_k)\left[1 - P_{\text{ext},B}(t_k,T)\right] \Big\}. \end{aligned} \tag{S5}$$

This expression accounts for the fact that single resistants can be of type $A$ or $B$, and that we are interested in situations where neither type is present. The terms $\overline{W}_A$ and $\overline{W}_B$ are the average production rates of mutants of type $A$ and $B$ from strain $S$, as defined in Eqs. (3) and (4) of the main text. To lowest order in $\Delta t$ the above expression turns into

$$P_{0,t_k}^{\text{single}}(T) \approx \exp\Big( - \Delta t \big\{ W_A(t_k)\left[1 - P_{\text{ext},A}(t_k,T)\right] + W_B(t_k)\left[1 - P_{\text{ext},B}(t_k,T)\right] \big\} \Big). \tag{S6}$$

The probability that no single resistant cells (of either type $A$ or type $B$) are present at time $T$ in turn takes the form

$$P_0^{\text{single}}(T) \equiv P\left(n_A(T) = n_B(T) = 0\right) = \prod_{k=1}^{N} P_{0,t_k}^{\text{single}}(T), \tag{S7}$$

since we require that single resistant cells produced in any interval $[t_k, t_k + \Delta t]$ get extinct along with their lineage. Inserting the expression from Eq. (S6) and taking the limit $\Delta t \to 0$ (so that sums turn into integrals), we find

$$P_R^{\text{single}}(T) = 1 - P_0^{\text{single}}(T) = 1 - \exp\left[ - \int_0^T \left\{ \Phi_A(t,T) + \Phi_B(t,T) \right\} \mathrm{d}t \right], \tag{S8}$$

with

$$\begin{aligned} \Phi_A(t,T) &= \overline{W}_A(t)\left[1 - P_{\text{ext},A}(t,T)\right], \\ \Phi_B(t,T) &= \overline{W}_B(t)\left[1 - P_{\text{ext},B}(t,T)\right]. \end{aligned} \tag{S9}$$

The calculation can be extended to the case of several single resistant types, $i = 1, \ldots, M$. One would get

$$P_R^{\text{single}}(T) = 1 - \exp\left[-\int_0^T \left(\sum_{i=1}^{M} \Phi_i\left(t,T\right)\right) \mathrm{d}t\right], \tag{S10}$$

with $\Phi_i$ analogous to $\Phi_A$ and $\Phi_B$ above.

##### S1.2.2 Double resistance

The procedure to obtain the probability of double resistance is similar to the above line of reasoning. The main difference is that there are now two sources, the two single resistant strains $A$ and $B$. As before, we start by writing down the probability that double resistant cells either do not arise in $[t_k, t_k + \Delta t]$, or, if they do, that they and their lineage goes extinct by time $T$. This probability is

$$P_{0,t_k}^{\text{double}}(T) = \left[1 - \overline{W}_D(t_k)\Delta t\right] + \overline{W}_D(t_k)\Delta t P_{\text{ext},D}(t_k, T), \tag{S11}$$

where $\overline{W}_D$ is the average production rate of mutants type $D$ (from strains $A$ or $B$), given in Eq. (5) in the main text. Following the same steps as before, the probability of having at least one double resistant cell at time $T$ becomes

$$P_R^{\text{double}}(T) = 1 - \exp\left[-\int_0^T \left\{\Psi_A\left(t,T\right) + \Psi_B\left(t,T\right)\right\} \mathrm{d}t\right], \tag{S12}$$

with

$$\begin{aligned} \Psi_A\left(t,T\right) &= b_A(t)\mu_B\bar{n}_A(t)\left[1 - P_{\text{ext},D}(t,T)\right], \\ \Psi_B\left(t,T\right) &= b_B(t)\mu_A\bar{n}_B(t)\left[1 - P_{\text{ext},D}(t,T)\right]. \end{aligned} \tag{S13}$$

We stress again that this approach is based on a deterministic approximation for the production rates $W_A, W_B$ and $W_D$. More specifically, these rates can – in principle – depend on the composition of the population, $\mathbf{n}$, which is a stochastic variable. In our approximation the components $n_i$ of $\mathbf{n}$ are replaced with their mean values $\bar{n}_i$, resulting for example in the appearance of $\bar{n}_A$ and $\bar{n}_B$ in Eq. (S13). If such an approximation is not made, then $W_A, W_B$ and $W_D$ are stochastic variables.

### S2 Probabilities of resistance for the exponential growth model (EG) with constant growth coefficients

The differential equations for the mean values of cell numbers for the exponential growth model (abbreviated 'EG' in the main paper) are given in Eq. (6) in the main text. For constant coefficients $b_i$ and $d_i$

they have the following analytical solutions

$$\begin{aligned}
\bar{n}_S(t) &= n_0 e^{r_S t}, \\
\bar{n}_A(t) &= \frac{n_0 b_S \mu_A}{r_S - r_A}\left(e^{r_S t} - e^{r_A t}\right), \\
\bar{n}_B(t) &= \frac{n_0 b_S \mu_B}{r_S - r_B}\left(e^{r_S t} - e^{r_B t}\right), \\
\bar{n}_D(t) &= n_0 b_S \mu_A \mu_B \times \\
&\left[\frac{1}{r_S - r_D}\left(\frac{b_A}{r_S - r_A} + \frac{b_B}{r_S - r_B}\right)\left(e^{r_S t} - e^{r_D t}\right) + \frac{b_A}{(r_S - r_A)(r_D - r_A)}\left(e^{r_A t} - e^{r_D t}\right)\right. \\
&\left. + \frac{b_B}{(r_S - r_B)(r_D - r_B)}\left(e^{r_B t} - e^{r_D t}\right)\right]. \qquad \text{(S14)}
\end{aligned}$$

If $b_A = b_B = 0$ (strains $A$ and $B$ do not replicate), then $\bar{n}_D(t) = 0$. Also, for $\mu_A = \mu_B = 0$ there are no single or double mutants as there are no mutations.

Moreover, for constant birth and death rates the expression in Eq. (S1) reduces to

$$P_{\text{ext},i}(t,T) = 1 - \frac{r_i}{b_i - d_i e^{-r_i(T-t)}}. \qquad \text{(S15)}$$

If $d_i = 0$, then $P_{\text{ext},i}(t,T) = 0$. We also note that the expression for $P_{\text{ext},i}(t,T)$ shows that as $T$ grows, two possible types of behaviour can occur at fixed $t$:

1. If $b_i > d_i$ ($r_i > 0$), then $P_{\text{ext},i}(t,T)$ tends to its maximum value $d_i/b_i$. The probability of extinction at long times is then less than one.
2. If $b_i < d_i$ ($r_i < 0$), then $P_{\text{ext},i}(t,T)$ tends to one as $T \to \infty$, i.e. extinction is certain in the long run.

For the simple case of constant birth and death rates model, it is also possible to derive analytical solutions for the probabilities of single and double resistance. For the former we get

$$P_R^{\text{single}}(T) = 1 - e^{-[\phi_A(T) + \phi_B(T)]}, \qquad \text{(S16)}$$

with

$$\phi_A(T) = \frac{n_0 b_S \mu_A r_A}{r_S b_A}\left[e^{r_S T}\, {}_2F_1\left(1, \frac{r_S}{r_A}; \frac{r_A + r_S}{r_A}; \frac{d_A}{b_A}\right) - {}_2F_1\left(1, \frac{r_S}{r_A}; \frac{r_A + r_S}{r_A}; \frac{d_A e^{-r_A T}}{b_A}\right)\right], \qquad \text{(S17)}$$

and

$$\phi_B(T) = \frac{n_0 b_S \mu_B r_B}{r_S b_B}\left[e^{r_S T}\, {}_2F_1\left(1, \frac{r_S}{r_B}; \frac{r_B + r_S}{r_B}; \frac{d_B}{b_B}\right) - {}_2F_1\left(1, \frac{r_S}{r_B}; \frac{r_B + r_S}{r_B}; \frac{d_B e^{-r_B T}}{b_B}\right)\right], \qquad \text{(S18)}$$

where ${}_2F_1$ is the Gaussian hypergeometric function (also known as the ordinary hypergeometric function). The terms $\phi_A$ and $\phi_B$ capture mutations from strain $S$ to $A$ or $B$, respectively. If at least one of the functions $\phi_A$ and $\phi_B$ tends to infinity as $T \to \infty$, then $P_R^{\text{single}}(T) \to 1$. This can happen if at least one of $r_A$ or $r_B$ is positive.

We also note that $P_R^{\text{single}}(T)$ is trivially zero if $n_0$ or $b_S$ vanish (no reproduction of sensitive cells), or if $\mu_A$ and $\mu_B$ are both zero (no mutations).

For constant birth and death rates $b_i$ and $d_i$ the probability of double resistance becomes

$$P_R^{\text{double}}(T) = 1 - e^{-[\psi_A(T) + \psi_B(T)]}, \qquad \text{(S19)}$$

where

$$\psi_A(T) = \frac{n_0 b_A \mu_B r_D b_S \mu_A}{b_D(r_S - r_A)} \left[\psi(r_S, r_D) - \psi(r_A, r_D)\right] \tag{S20}$$

and

$$\psi_B(T) = \frac{n_0 b_B \mu_B r_D b_S \mu_A}{b_D(r_S - r_B)} \left[\psi(r_S, r_D) - \psi(r_B, r_D)\right]. \tag{S21}$$

We have defined

$$\psi(r_l, r_D) = \frac{1}{r_\ell} \left[ e^{r_\ell T} {}_2F_1\left(1, \frac{r_\ell}{r_D}; \frac{r_\ell + r_D}{r_D}; \frac{d_D}{b_D}\right) - {}_2F_1\left(1, \frac{r_\ell}{r_D}; \frac{r_\ell + r_D}{r_D}; \frac{d_D e^{-r_D T}}{b_D}\right)\right], \tag{S22}$$

for $\ell = S, A, B$. The terms $\psi_A$ and $\psi_B$ take into account contributions from strains $A$ and $B$ that mutate into strain $D$, respectively. If $n_0$ or $b_S$ vanish (no growth of sensitive cells), or $\mu_A$ and $\mu_B$ are zero (no mutations), or both $b_A$ or $b_B$ are zero (no growth of single resistant strains), then $P_R^{\text{double}}(T) = 0$, i.e., double resistance does not emerge.

### S3 Probabilities of resistance for the logistic growth model without competition between strains (LG)

#### S3.1 Constant coefficients

##### S3.1.1 Mean number of sensitive cells

For constant drug concentrations, the functions $d_i(t)$ (death rate) and $\rho_i(t)$ (intrinsic growth rate) are constant in time. The birth rate $b_i(t)$, however, depends on $n_i(t)$, so it is not constant [see Eq. (7) of the main paper].

For this model, there is no analytical solution to the differential equations for the mean cell numbers [Eqs. (1) in the main paper], except for strain $S$, for which one finds the logistic function

$$\bar{n}_S(t) = \frac{k_S}{1 + (k_S/n_0 - 1)\, e^{-\rho_S t}}. \tag{S23}$$

For the other strains, it is necessary to integrate the equations for the mean cell numbers numerically.

##### S3.1.2 Extinction probability for a single-species logistic birth-death process

In this section we derive a closed-form solution for the probability of extinction, $P_{\text{ext},i}$ of a single-species logistic birth-death process with time dependent rates. To do this, we follow a procedure similar to the one in [4] for the Gompertz model.

We focus on strain $i$, and start from the birth rate

$$b_i(t) = d_i + \rho_i \left(1 - \frac{\bar{n}_i(t)}{k_i}\right), \tag{S24}$$

in which we have made the deterministic approximation, i.e., we have replaced the number of cells of type $i$ by its mean value. This turns the birth rate into a time-dependent external function. The death rate $d_i$ is constant, as is the rate $\rho_i$. However, the actual births and deaths are treated as stochastic events.

We label time by $t'$ and assume that at a certain time $t' = t$ there is only one individual, $\bar{n}_i(t' = t) = 1$.

The function $\bar{n}_i(t')$ follows the differential equation

$$\frac{\mathrm{d}\bar{n}_i}{\mathrm{d}t'}(t') = \rho_i \left(1 - \frac{\bar{n}_i(t')}{k_i}\right) \bar{n}_i(t'). \tag{S25}$$

The solution of this equation, subject to the condition $\bar{n}_i(t'=t) = 1$, is

$$\bar{n}_i(t') = \frac{k_i}{1 + (k_i - 1)\, e^{-\rho_i(t'-t)}}. \tag{S26}$$

Then, using Eq. (19), we write

$$\beta_i\,(t, t') = \log\, \bar{n}_i(t'), \tag{S27}$$

so that,

$$\begin{aligned}\int_t^T d_i e^{-\beta_i(t,t')} \mathrm{d}t' &= d_i \int_t^T \frac{1}{\bar{n}_i(t')} \mathrm{d}t' \\ &= \frac{d_i}{k_i}\left[(T-t) - \frac{(k_i-1)}{r_i}\left(e^{-\rho_i(T-t)-1}\right)\right].\end{aligned} \tag{S28}$$

Finally, putting all together in Eq. (S1), we obtain

$$P_{\mathrm{ext},i}(t,T) = \frac{\dfrac{d_i}{k_i}\left[(T-t) - \dfrac{(k_i-1)}{\rho_i}\left(e^{-\rho_i(T-t)} - 1\right)\right]}{\dfrac{d_i}{k_i}\left[(T-t) - \dfrac{(k_i-1)}{\rho_i}\left(e^{-\rho_i(T-t)} - 1\right)\right] + 1}. \tag{S29}$$

This is the extinction probability for a single individual and its lineage in a single-species logistic growth model with constant coefficients $\rho_i$ and $d_i$. Trivially, $P_{\mathrm{ext},i}(t,T) = 0$ for $d_i = 0$.

##### S3.1.3 Single resistance

For the logistic model without competition between strains (LG), the average production rates of strains $A$ and $B$ are given by

$$\overline{W}_A(t) = \left[d_S + \rho_S\left(1 - \frac{\bar{n}_S(t)}{k_S}\right)\right] \mu_A \bar{n}_S(t) \tag{S30}$$

$$\overline{W}_B(t) = \left[d_S + \rho_S\left(1 - \frac{\bar{n}_S(t)}{k_S}\right)\right] \mu_B \bar{n}_S(t). \tag{S31}$$

We focus again on the case of constant coefficients $d_i$ and $\rho_i$. Using the closed-form solution in Eq. (S29) for the extinction probability, we use Eq. (S9) to write

$$\Phi_A\,(t,T) = \frac{\left[d_S + \rho_S\left(1 - \dfrac{\bar{n}_S(t)}{k_S}\right)\right] \mu_A \bar{n}_S(t)}{\dfrac{d_A}{k_A}\left[(T-t) - \dfrac{(k_A-1)}{\rho_A}\left(e^{-\rho_A(T-t)} - 1\right)\right] + 1}, \tag{S32}$$

and

$$\Phi_B\,(t,T) = \frac{\left[d_S + \rho_S\left(1 - \dfrac{\bar{n}_S(t)}{k_S}\right)\right] \mu_B \bar{n}_S(t)}{\dfrac{d_B}{k_B}\left[(T-t) - \dfrac{(k_B-1)}{\rho_B}\left(e^{-\rho_B(T-t)} - 1\right)\right] + 1}. \tag{S33}$$

In these expressions, $\bar{n}_S$ is a logistic function with initial condition $\bar{n}_S(t=0) = n_0$ and growth rate $\rho_S$, i.e. $\bar{n}_S(t) = k_S/\left(1+(k_S/n_0)e^{-\rho_S t}\right)$.

The expressions for $\Phi_A$ and $\Phi_B$ are integrated with respect to $t$ in Eq. (S8). Even though we can write down $\Phi_A$ and $\Phi_B$ in the above form, we have not been able to find an analytical solution for these integrals. The probability of single resistance is obtained by performing these integrals numerically.

###### S3.1.4 Double resistance

For constant coefficients, the production rate of strain $D$ in Eq. (5) of the main paper takes the form

$$\overline{W}_D(t) = \left[d_A + \rho_A\left(1-\frac{\bar{n}_A(t)}{k_A}\right)\right]\mu_B\bar{n}_A(t) + \left[d_B + \rho_B\left(1-\frac{\bar{n}_B(t)}{k_B}\right)\right]\mu_A\bar{n}_B(t), \tag{S34}$$

for the logistic growth model with no competition between strains. Eqs. (17) and (18) become

$$\Psi_A(t,T) = \frac{\left[d_A + \rho_A\left(1-\frac{\bar{n}_A(t)}{k_A}\right)\right]\mu_B\bar{n}_A(t)}{\frac{d_D f_D}{k_D}\left[(T-t) - \frac{(k_D-1)}{\rho_D}\left(e^{-\rho_D(T-t)}-1\right)\right]+1}, \tag{S35}$$

and

$$\Psi_B(t,T) = \frac{\left[d_B + \rho_B\left(1-\frac{\bar{n}_B(t)}{k_B}\right)\right]\mu_A\bar{n}_B(t)}{\frac{d_D f_D}{k_D}\left[(T-t) - \frac{(k_D-1)}{\rho_D}\left(e^{-\rho_D(T-t)}-1\right)\right]+1}. \tag{S36}$$

While the function $\bar{n}_S(t)$ is available in closed form [see Eq. (S23)], no analytical solutions were found for $\bar{n}_A(t)$ and $\bar{n}_B(t)$. This is because the single resistant types are produced from $S$, and $\bar{n}_S$ itself is a time-dependent quantity. The functions $\bar{n}_A(t)$ and $\bar{n}_B(t)$ are therefore obtained by numerical integration of the differential equations (1) in the main paper.

##### S3.2 Time-dependent rates

We now assume that the rates $d_i(t)$ and $\rho_i(t)$ in Eq. (7) of the main text are functions of time. In order to obtain the extinction probability in an effective way, we use Eq. (S27) to write Eq. (S1) as

$$P_{\text{ext},i}(t,T) = \frac{\int_t^T \frac{d_i(t')}{\bar{n}_i(t')}\mathrm{d}t'}{1+\int_t^T \frac{d_i(t')}{\bar{n}_i(t')}\mathrm{d}t'}, \tag{S37}$$

with $\bar{n}_i(t')$ as in Eq. (20), see also Sec. 3.4 of the main paper.

To derive Eq. (20), i.e. to express the solutions for $\bar{n}_i(t')$ (with initial condition $\bar{n}_i(t) = 1$) in terms of solutions $\bar{n}_i^0(t')$ (with $\bar{n}_i^0(0) = 0$), we integrate Eq. (S25) to obtain

$$\bar{n}_i(t') = \frac{k_i}{(k_i-1)e^{-\int_t^{t'}\rho_i(t'')\mathrm{d}t''}+1}. \tag{S38}$$

Similarly,

$$\bar{n}_i^0(t') = \frac{k_i}{(k_i-1)e^{-\int_0^{t'}\rho_i(t'')\mathrm{d}t''}+1}, \tag{S39}$$

such that $\bar{n}_i^0(0) = 1$. From this, we get

$$e^{-\int_0^{t'} \rho_i(t'')\mathrm{d}t''} = \frac{k_i/\bar{n}_i^0(t') - 1}{k_i - 1}, \tag{S40}$$

and then

$$e^{-\int_t^{t'} \rho_i(t'')\mathrm{d}t''} = \frac{k_i/\bar{n}_i^0(t') - 1}{k_i/\bar{n}_i^0(t) - 1}. \tag{S41}$$

Inserting this result into Eq. (S38) gives Eq. (20) in the main text, i.e.,

$$\bar{n}_i(t' \geq t) = \frac{k_i\left(\frac{k_i}{\bar{n}_i^0(t)} - 1\right)}{(k_i - 1)\left(\frac{k_i}{\bar{n}_i^0(t')} - 1\right) + \left(\frac{k_i}{\bar{n}_i^0(t)} - 1\right)}. \tag{S42}$$

The probabilities of resistance are then obtained by performing the integrals in Eqs. (S8) and (S12) numerically.

#### S4 Probabilities of resistance for the logistic model with between-strain competition (CLG)

As explained in Section 4.1 of the main paper, we have not been able to derive analytical solutions of the mean cell numbers for the logistic growth model with competition between the strains. This is due to the coupling of the equations for the different $\bar{n}_i$, see Eq. (1) in the main paper, and the rates in Eq. (8). Because of the lack of closed form expressions for the $\bar{n}_i$ we cannot find an analytical solution for the extinction probability $P_{\mathrm{ext},i}$ either, i.e., there is no closed-form equivalent of Eq. (S29) in the CLG model, even when the coefficients $d_i$ and $\rho_i$ are constant.

As a consequence the $\bar{n}_i(t)$ are obtained from numerical integration of the differential equations for the mean cell numbers [Eq. (1)]. The extinction probabilities are then obtained from Eq. (S37), where the required integral is again evaluated numerically. The integration is carried out by applying the strategy described in Section 3.4 of the main paper. We detail below how to apply it for the CLG model. This procedure is used both for constant and time-dependent coefficients $d_i$ and $\rho_i$.

In order to calculate $P_{\mathrm{ext},i}$, we focus on a single species following a birth-death process with birth rate

$$b_i(t) = \begin{cases} d_i(t) + \rho_i(t)\left(1 - \frac{\bar{n}_T(t)}{k_i}\right) & \text{if} \quad \bar{n}_T(t) \leq k_i \\ d_i(t) & \text{if} \quad \bar{n}_T(t) > k_i, \end{cases} \tag{S43}$$

where $d_i(t)$ is a given death rate, and $\rho_i(t)$ the intrinsic birth rate of species $i$. As explained in Section 3.4, to obtain the extinction probability in an efficient manner we need to express solutions $\bar{n}_i(t')$ with initial condition $\bar{n}_i(t' = t) = 1$ in terms of solutions $\bar{n}_i^0(t')$ that satisfy $\bar{n}_i^0(t' = 0) = 1$. Notice that, although the focus is on species $i$, we cannot ignore the effect of the remaining species for the model with competition between strains: the function $\bar{n}_T(t')$ in Eq. (S43) accounts for the mean value of the total number of cell across all species. The function $\bar{n}_i(t')$ follows

$$\frac{\mathrm{d}\bar{n}_i}{\mathrm{d}t'}(t') = \rho_i(t')\left(1 - \frac{\bar{n}_T(t')}{k_i}\right)\bar{n}_i(t'). \tag{S44}$$

To proceed, we treat $\bar{n}_T(t')$ on the right hand-side of Eq. (S44) as a given function. Formally integrating we then obtain

$$\bar{n}_i(t') = \exp\left\{\int_t^{t'} \rho_i(t'')\left(1 - \frac{\bar{n}_T(t'')}{k_i}\right) \mathrm{d}t''\right\}, \tag{S45}$$

where we note the initial condition $\bar{n}_i(t' = t) = 1$ The solution $\bar{n}_i^0(t')$ with condition $\bar{n}_i^0(0) = 0$ (see Section 3.4 of the main paper) is given by

$$\bar{n}_i^0(t') = \exp\left\{\int_0^{t'} \rho_i(t'')\left(1 - \frac{\bar{n}_T(t'')}{k_i}\right) \mathrm{d}t''\right\}. \tag{S46}$$

Using these one obtains

$$\bar{n}_i(t' \geq t) = \frac{\bar{n}_i^0(t')}{\bar{n}_i^0(t)}, \tag{S47}$$

with which $P_{\mathrm{ext},i}$ is obtained from Eq. (S37) and from it, the probabilities of single and double resistance from Eqs. (S8) and (S12).

### S5 Further supplementary results

In this section, we present additional data from simulations and from the analytical approach. These complement the results in the main paper.

#### S5.1 Probability of double resistance for constant growth rates

Figure S1 complements Fig. 4. It shows how the probability of double resistance, $P_R^{\text{double}}$, differs across the different growth models as a function of parameters not varied in Fig. 4. As in Fig. 4 parameters are varied in turn, i.e., all but the one shown on the vertical axis are kept fixed.

The data in the figure shows that varying $d_A$ the emergence of double resistance shows a noticeable delay in the CLG model compared to the other two growth models. When only $\mu_A$ varies, however, all the models have similar probability of double resistance. This is because $\mu_B$ is high enough that double resistance does not get affected. This example shows that competition may not significantly affect the emergence of double resistance in some circumstances.

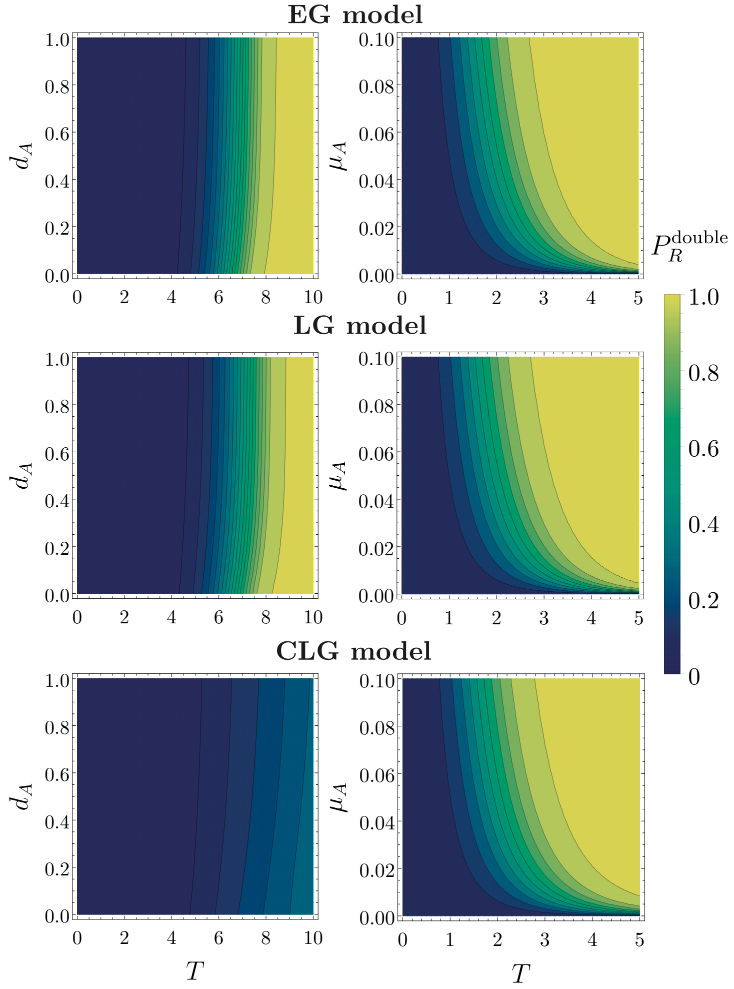


**Figure S1:** Theoretical prediction of the probability of double resistance for the three growth models when varying one parameter. When not varied, we use the same parameters as in Figure 4 of the main text, i.e. $b_S = 1.0, b_A = 1.1, b_B = 1.2, b_D = 1.3, d_S = d_A = d_B = d_D = 0.1, \mu_A = \mu_B = 10^{-4}$, $n_0 = 10^4$, $k_S = 10^6, k_A = 1.1 \times 10^6, k_B = 1.2 \times 10^6$, and $k_D = 1.3 \times 10^6$.

#### S5.2 Further tests of theoretical predictions for resistance against numerical simulations

In Figs. S2–S4 we show data similar to that in Fig. 1 in the main paper, but for different choices of the model parameters. The figure shows the probabilities of single and double resistance as a function of time, both from simulations and as predicted from the theory. The parameters in Figs. S2–S4 are chosen such that they produce a noticeable difference in the prediction of double resistance relative to the results in Fig. 1, in particular for the model with competition between strains. We do this based on the results presented in Fig. 4 of the main paper, i.e., we select parameter values such that the delay in the emergence of double resistants is noticeably higher than with the parameters from Fig. 1.

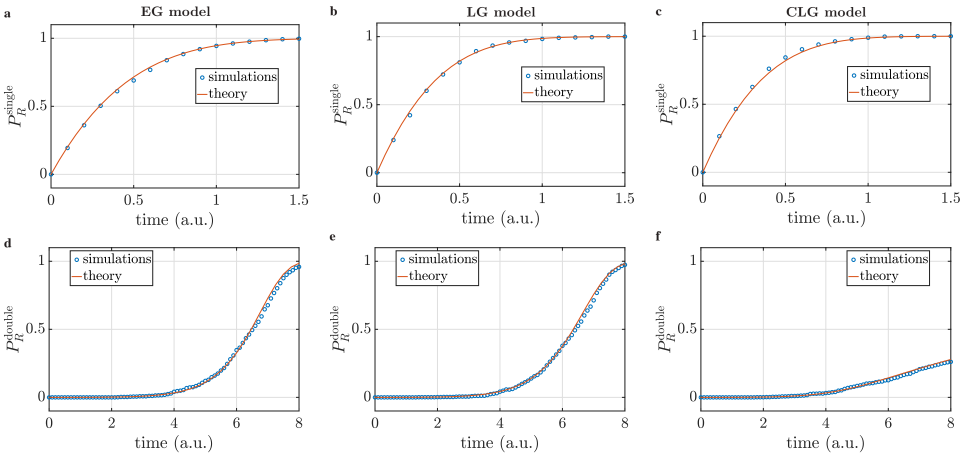


**Figure S2:** Probabilities of single and double resistance for the same parameters as in Figure 4 of the main text but with $d_S = 0.5$.

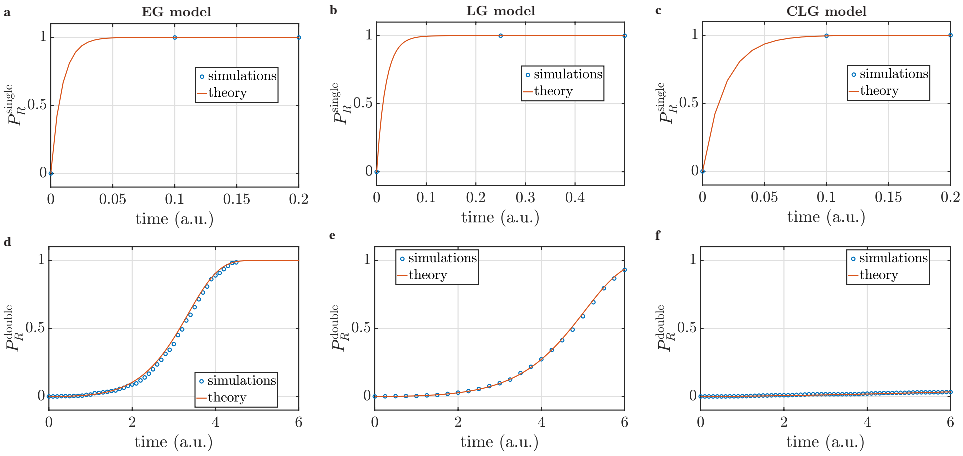


**Figure S3:** Probabilities of single and double resistance for the same parameters as in Figure 4 of the main text but with $n_0 = 5 \times 10^5$.

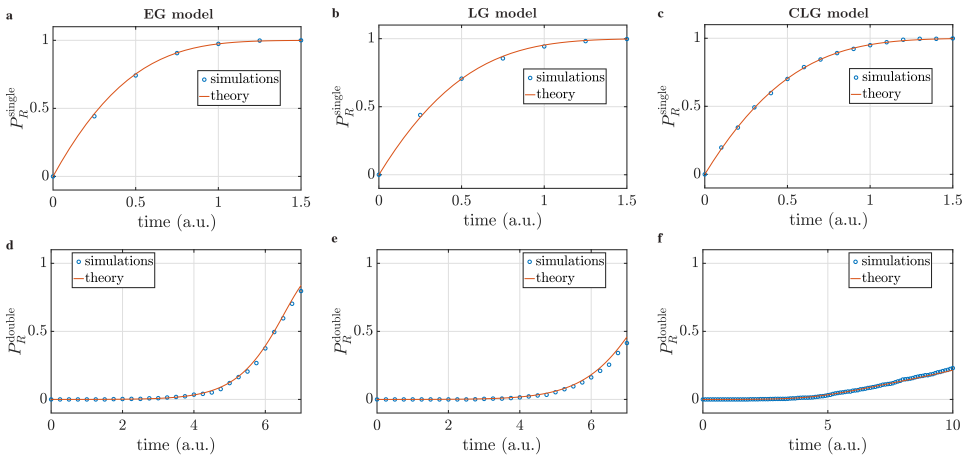


**Figure S4:** Probabilities of single and double resistance for same parameters as in Figure 4 of the main text but with $r_A = 0.1$.

#### S5.3 Probability of single and double resistance for time-dependent dosing schedules

In this section, we complement the results for the probabilities of single and double resistance shown in Section 7.

In Fig. S5, we show the theoretical prediction for the probability of single resistance using the same parameters and dosing schedules (sinusoidal and pulsing therapies) as in Fig. 7 of the main paper. As shown, the probability of single resistance is noticeably different in the exponential model than for the two models with logistic growth. Therefore it is important to consider carefully the choice of growth model to best describe a given biological system. If an exponential model is used, but the real-world system is subject to restricted growth then the prediction for example of the typical time at which first resistants emerge may not be accurate. Further, the value of $\Delta t_B$ that optimises the treatment (most delays the emergence of single resistance) and the typical time at which the first single mutants emerge are both different in the exponential growth model than in the two logistic models.

In Figures S6 and S7, we re-plot data shown in Fig. 7 of the main paper. We now illustrate more clearly the effect of changing $\Delta t_B$ in each growth model. Figure S6 demonstrates that the emergence of double resistance is considerably delayed by competition. In Fig. S7 we show the probability of double resistance, $P_R^{\text{double}}(T)$, as function of $\Delta t_B$ for a given time $T$. This is shown for each of the three growth models, and for the sinusoidal and pulsing periodic treatments. The figure demonstrates that the optimum treatment is different for each of the three growth models.

In Fig. S8 finally we show the probability of double resistance with a higher initial sensitive cell number and higher carrying capacities values for each strain than in Fig. 7. For these parameters, the prediction of the competition model shows a more pronounced difference than the prediction in Fig. 7 when varying $\Delta t_B$.

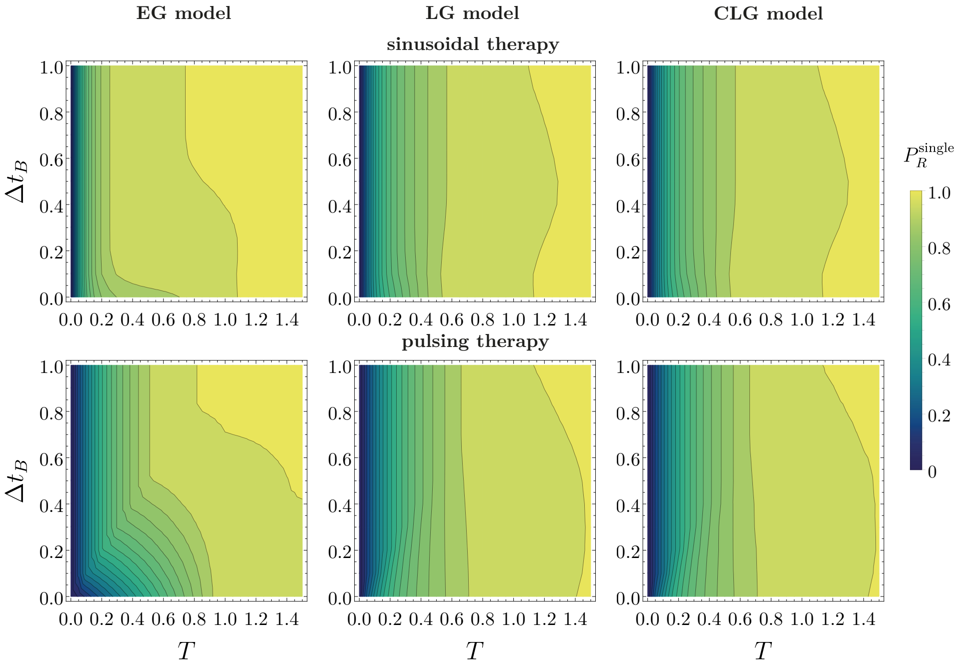


**Figure S5:** Theoretical prediction of single resistance for sinusoidal and pulsing drug therapies for parameters of Figure 7 from the main text: $b_S = b_A = b_B = b_D = 1.1, d_S = d_A = d_B = d_D = 0.1, \mu_A = \mu_B = 10^{-3}$, $n_0 = 5 \times 10^3$, $k_S = 10^4, k_A = 1.1 \times 10^4, k_B = 1.2 \times 10^4$, and $k_D = 1.3 \times 10^4$.

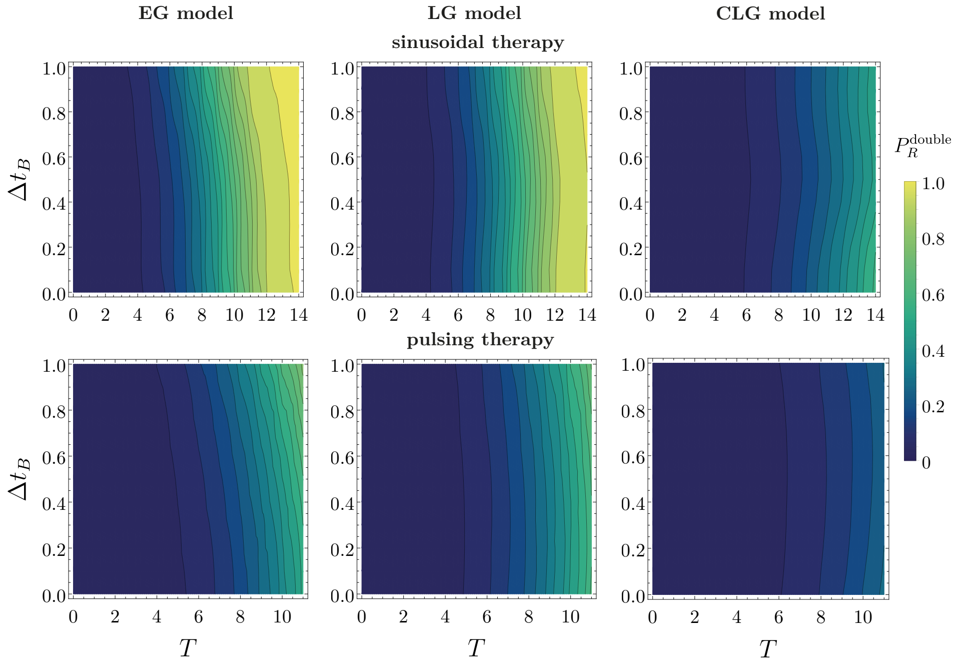


**Figure S6:** Theoretical prediction of the probability of double resistance for sinusoidal and pulsing drug therapies for parameters of Figure 7 from the main text: $b_S = b_A = b_B = b_D = 1.1, d_S = d_A = d_B = d_D = 0.1, \mu_A = \mu_B = 10^{-3}$, $n_0 = 5 \times 10^3$, $k_S = 10^4, k_A = 1.1 \times 10^4, k_B = 1.2 \times 10^4$, and $k_D = 1.3 \times 10^4$

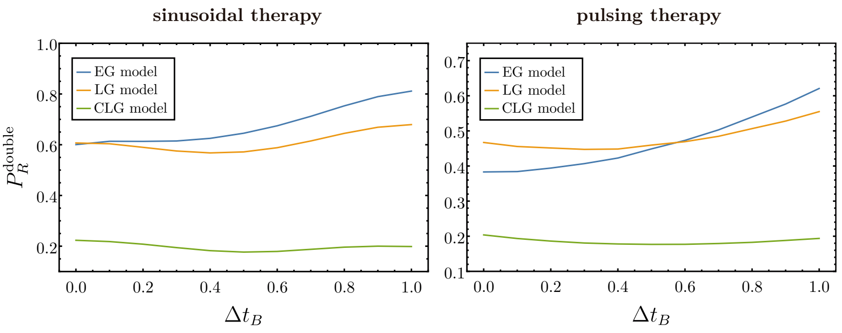


**Figure S7:** Theoretical prediction of the probability of double resistance for each growth model during sinusoidal (left panel) and pulsing therapy (right panel) as a function of $\Delta t_B$ at time $T = 10$ after the treatment started. Parameters used are the same in Figure 7 from the main text: $b_S = b_A = b_B = b_D = 1.1, d_S = d_A = d_B = d_D = 0.1, \mu_A = \mu_B = 10^{-3}$, $n_0 = 5 \times 10^3$, $k_S = 10^4, k_A = 1.1 \times 10^4, k_B = 1.2 \times 10^4$, and $k_D = 1.3 \times 10^4$.

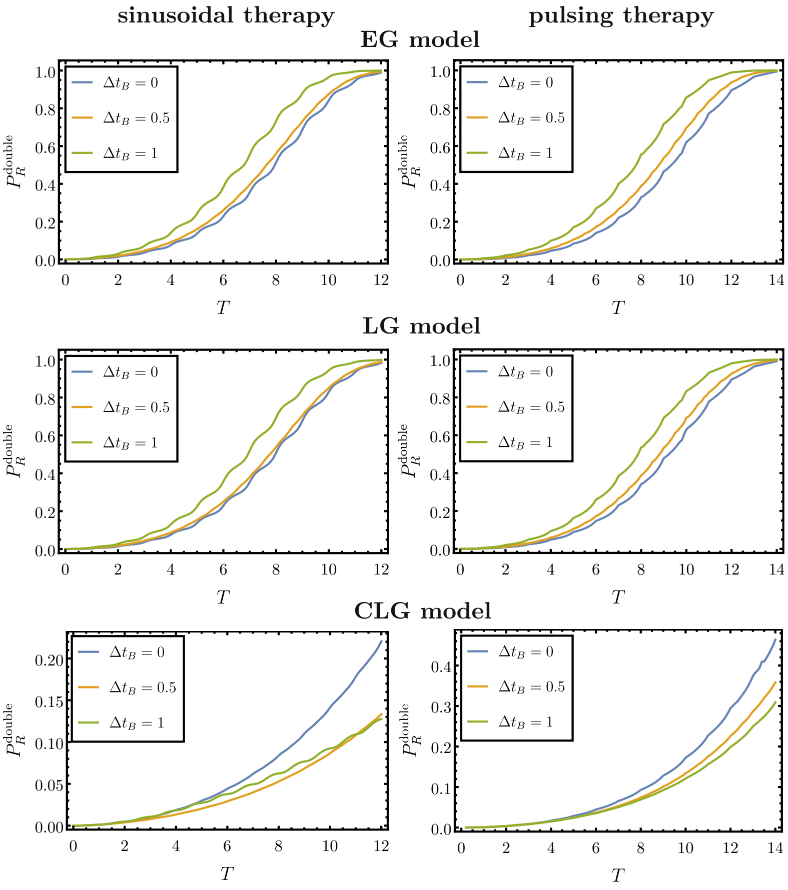


**Figure S8:** Theoretical prediction of the probability of double resistance for each growth model when varying $\Delta t_B$ for sinusoidal and pulsing therapies. Parameters used: $b_S = b_A = b_B = b_D = 1.1, d_S = d_A = d_B = d_D = 0.1, \mu_A = \mu_B = 10^{-3}$, $n_0 = 10^4$, $k_S = 10^5, k_A = 1.1 \times 10^4, k_B = 1.2 \times 10^4$, and $k_D = 1.3 \times 10^4$.

### S6 Limitations of the analytical approach

In Fig. S9, we show an instance in which the theoretical predictions for the probability of double resistance deviate from results from simulations. Specifically, we have chosen smaller birth rates for each strain in panel (a), and smaller mutation rates in panels (b) and (c) than in Fig. 1 of the main paper. In either case, the overall effect is a delay in the emergence of double resistants. As explained in Section 6 the theoretical predictions then become less accurate.

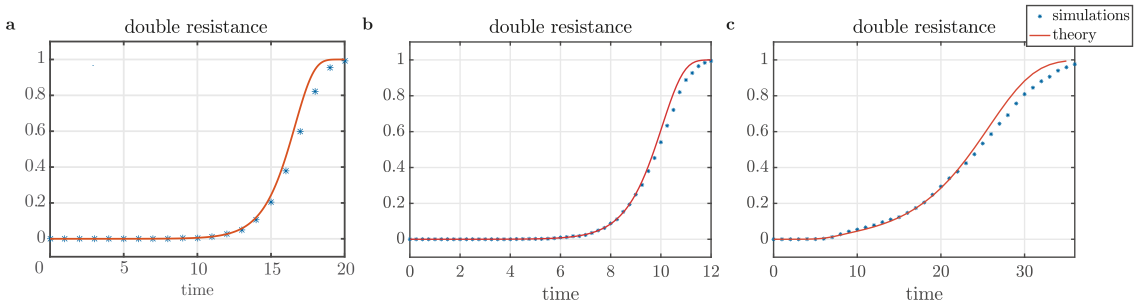


**Figure S9:** Theoretical prediction of double resistance for: (a) Exponential growth model for parameters: $b_S = 0.5, b_A = 0.8, b_B = 0.9, b_D = 1.0, d_S = d_A = d_B = d_D = 0.1, \mu_A = 10^{-6}, \mu_B = 10^{-4}$, $n_0 = 10^4$. (b) Non-competitive growth model for parameters: $b_S = 1.1, b_A = 1.2, b_B = 1.3, b_D = 1.4, d_S = d_A = d_B = d_D = 0.1, \mu_A = \mu_B = 10^{-5}, k_S = 10^7, k_A = 1.1 \times 10^7, k_B = 1.2 \times 10^7$, and $k_D = 1.3 \times 10^7$. (c) Competitive growth model for parameters: $b_S = 1.1, b_A = 1.2, b_B = 1.3, b_D = 1.4, d_S = d_A = d_B = d_D = 0.1, \mu_A = \mu_B = 10^{-5}, k_S = 10^7, k_A = 1.1 \times 10^7, k_B = 1.2 \times 10^7$, and $k_D = 1.3 \times 10^7$.

[3] Emanuel Parzen. *Stochastic processes.* SIAM, San Francisco, 1999.

[4] PR Parthasarathy and B Krishna Kumar. A birth and death process with logistic mean population. *Communications in Statistics-Theory and Methods*, 20(2):621–629, 1991.
